# Whole brain imaging reveals distinct spatial patterns of amyloid beta deposition in three mouse models of Alzheimer’s disease

**DOI:** 10.1101/395236

**Authors:** Jennifer D. Whitesell, Alex R. Buckley, Joseph E. Knox, Leonard Kuan, Nile Graddis, Andrew Pelos, Alice Mukora, Wayne Wakeman, Phillip Bohn, Anh Ho, Karla E. Hirokawa, Julie A. Harris

## Abstract

A variety of Alzheimer’s disease (AD) mouse models overexpress mutant forms of human amyloid precursor protein (APP), producing high levels of amyloid β (Aβ) and forming plaques However, the degree to which these models mimic spatiotemporal patterns of Aβ deposition in brains of AD patients is unknown. Here, we mapped the spatial distribution of Aβ plaques across ages in three APP-overexpression mouse lines (APP/PS1, Tg2576, hAPP-J20) using *in vivo* labeling with methoxy-X04, high throughput whole brain imaging, and an automated informatics pipeline. Images were acquired with high resolution serial 2-photon tomography and labeled plaques were detected using custom-built segmentation algorithms. Image series were registered to the Allen Mouse Brain Common Coordinate Framework, a 3D reference atlas, enabling automated brain-wide quantification of plaque density, number, and location. In both APP/PS1 and Tg2576 mice, plaques were identified first in isocortex, followed by olfactory, hippocampal, and cortical subplate areas. In hAPP-J20 mice, plaque density was highest in hippocampal areas, followed by isocortex, with little to no involvement of olfactory or cortical subplate areas. Within the major brain divisions, distinct regions were identified with high (or low) plaque accumulation; *e.g.*, the lateral visual area within the isocortex of APP/PS1 mice had relatively higher plaque density compared with other cortical areas, while in hAPP-J20 mice, plaques were densest in the ventral retrosplenial cortex. In summary, we show how whole brain imaging of amyloid pathology in mice reveals the extent to which a given model recapitulates the regional Aβ deposition patterns described in AD.

## 1. Introduction

Alzheimer’s disease (AD) is classically defined after death by the presence of two neuropathologies, amyloid β (Aβ) plaques and tau tangles. Recently, widespread advances in the use of biomarkers as rigorous measures of these pathologies in living people has led to a coordinated proposal for new definitions and staging of AD, incorporating biomarker profiles specific for Aβ and tau (Jack et al., 2018). One major benefit of using biomarkers to define disease stages is the ability to then design experiments testing novel therapeutics with the goal of intervening at early, presymptomatic stages. This will likely accelerate translational efforts, but does not replace the need for animal models early in the discovery and testing process. Thus, an important goal for the preclinical research field is to systematically characterize whether existing and novel mouse models adequately mimic the distribution and progression of these pathologies as mapped in human patients.

Pathological alterations in Aβ are currently the earliest detectable biomarker change, occurring before changes in tau pathologies, and sometimes decades before clinical symptoms (Jack et al., 2013). Most commonly used mouse models were engineered to express mutant forms of the amyloid precursor protein (APP), and/or presenilin 1 (PS1) which cause early onset forms of AD. These mutant APP mouse lines develop amyloid pathology, but little to no tau pathology or frank neurodegeneration, suggesting that they might best model early stages of AD (Sasaguri et al., 2017). Although mouse models have received much of the blame for the lack of success in translating preclinical research findings to approved therapeutics for clinical use, one possibility may be that these failures are due to misalignments in the disease stage modeled by these mice compared to the stage at which interventions are made in the clinical trials.

The large variety, accessibility, and sheer number of Alzheimer’s disease mouse models is an immensely important resource, revealing many novel basic science insights into Aβ pathologies. However, characterization of these lines with respect to molecular, circuit, network, and cognitive alterations is still very much incomplete, although at least one good resource based on post-hoc compilation of results exists (e.g., https://www.alzforum.org/research-models). As these data come from many independent labs using only a single line, often with different experimental focus on selected brain areas, and different methods and techniques, it is difficult to compare results across lines or interpret reported differences. Systematic characterization of mouse models, and a centralized database of results, would be a large asset for the AD community, assisting researchers in selecting the most appropriate lines based on experimental needs. New large-scale collaborative efforts promise to make progress toward this goal, at least for newly developed mouse models (e.g., MODEL-AD, https://model-ad.org/).

Here, our goal was to demonstrate the use of systematic whole brain imaging to characterize key pathological features in multiple mouse lines. We modified a high-throughput imaging and informatics pipeline, first developed for our Allen Mouse Connectivity Atlas project (Oh et al., 2014; http://connectivity.brain-map.org/projection), to label and map regional progression patterns of Aβ plaques across the entire brain in three frequently used APP mouse models: APP/PS1 (Jankowsky et al., 2001), hAPP-J20 (Mucke et al., 2000) and Tg2576 (Hsiao et al., 1996). Plaques were labeled across the entire brain via systemic injections of methoxy-X04, a fluorescent Congo red derivative that crosses the blood-brain barrier (Klunk et al., 2002). Previous reports of plaque density in these AD mouse models report different ages of onset (Hall & Roberson, 2012; J.-E. Lee & Han, 2013), with numerous reports of plaque density in cortex and hippocampus (Garcia-Alloza et al., 2006; H. Huang et al., 2016; Jährling et al., 2015; E. B. Lee et al., 2006; Liu et al., 2017; Mucke et al., 2000; Samaroo et al., 2012; Wright et al., 2013; Zhang et al., 2017). Some recent studies have reported plaque density for large subdivisions of the cortex (Kim et al., 2012) or a subset of structures (Liebmann et al., 2016) but they do not comprehensively describe plaque loads within subregions of these major brain divisions or across the rest of the brain.

In humans, progression and regional patterns of amyloid pathology have been described based on autopsy cases (Thal, Rüb, Orantes, & Braak, 2002), but also more increasingly with amyloid PET imaging (Buckner, 2005; Grothe et al., 2017; Rice & Bisdas, 2017). From these studies, we know that Aβ deposition occurs selectively first in the cortex, followed by hippocampal regions, including entorhinal and CA1, then other subcortical regions; e.g., striatum, basal forebrain, thalamus, and finally brainstem nuclei and cerebellum. Within the cortex, Aβ aggregates appear first, and are heaviest, in associational cortical areas, and specifically in the default mode network (DMN), which includes the precuneus, posterior cingulate cortex (PCC), retrosplenial cortex (RSP), medial prefrontal cortex, and lateral posterior parietal cortex (Buckner, Andrews-Hanna, & Schacter, 2008; Raichle et al., 2001).

Our results show plaque load is densest and appears earliest in the isocortex in both APP/PS1 and Tg2576 mice, like the early amyloid phases described by Thal et al., (2002). In contrast, plaque density was highest in hippocampal areas first, followed by isocortex in hAPP-J20 mice. We also identified plaques in select subcortical structures, mostly in the APP/PS1 line, in areas homologous to those described in the later amyloid deposition phases. Within the isocortex, the hAPP-J20 mice appeared to more closely mimic early stage human AD regional amyloid deposition; plaque load was higher in associational cortical areas as opposed to sensory and motor regions. Thus, systematic whole brain imaging of amyloid pathology in mice reveals line-specific regional deposition patterns. These data can be used together with characterization of other pathologies to identify the most suitable mouse models for testing early interventions in the progression of Alzheimer’s disease.

## 2. Materials and Methods

### Animals

All experimental procedures related to the use of mice were approved by the Institutional Animal Care and Use Committee of the Allen Institute for Brain Science, in accordance with NIH guidelines. We used heterozygous APP^+/-^ mice from the following transgenic lines: ***APP/PS1*** (B6.Cg-Tg(APPswe,PSEN1dE9)85Dbo/Mmjax, MMRRC Stock No: 034832-JAX) (Jankowsky et al., 2001)***, hAPP-J20*** (B6.Cg-Zbtb20Tg(PDGFB-APPSwInd)20Lms/2Mmjax, MMRRC Stock No: 34836-JAX) (Mucke et al., 2000), ***Tg2576*** (B6;SJL-Tg(APPSWE)2576Kha) (Hsiao et al., 1996). All animals were group-housed with a 10/14 light cycle (lights on from 6 AM to 8 PM, temperature = 68-72 degrees, humidity = 30-70%). APP/PS1 and hAPP-J20 mice were on the C57Bl/6J background and Tg2576 mice were on an FVB background. Mice were separated into six groups by age at perfusion: 5 months (P141-P156), 7 months (P202-P218), 9 months (P263-P307), 13 months (P386-P423), 19 months (P529-P589). The number of mice from each sex in each age group/transgenic line combination is listed in **Table 1**. We only observed very minor differences between the sexes in one region that had very low plaque density (the thalamus in hAPP-J20 mice), so we pooled male and female brains for all analyses (however, the two sexes were not equally distributed in our dataset, see **Table 1**). Our control dataset contained 35 nontransgenic (APP^−/−^) littermates from 7 - 19 months old from the APP/PS1 and hAPP-J20 lines (15 APP/PS1, 20 hAPP-J20; details in **Table 1**). All mice used in this study received a stereotaxic injection of AAV2/1.pCAG.FLEX.EGFP in the left hemisphere 20-25 days before perfusion; analyses of these data are not included in the current study. Informatics processing including segmentation and registration were performed on whole brain images, but all quantification was performed in the right hemisphere to minimize potential interference from the stereotaxic injection or EGFP fluorescence on plaque measurements.

**Table 1.** Experimental Dataset.

| <u>Mouse Line</u> | <u>Promoter</u> | <u>5 mo</u><br><u>(M/F)</u> | <u>7 mo</u><br><u>(M/F)</u> | <u>9 mo</u><br><u>(M/F)</u> | <u>13 mo</u><br><u>(M/F)</u> | <u>19 mo</u><br><u>(M/F)</u> |
| --- | --- | --- | --- | --- | --- | --- |
| APP/PS1 (APP<sup>+/-</sup>) | moPrP | 6 (4/2) | 13 (9/4) | 6 (3/3) | 39 (36/3) | 9 (7/2) |
| hAPP-J20 (APP<sup>+/-</sup>) | PDGF-β | 4 (2/2) | 3 (2/1) | 5 (2/3) | 11 (6/5) | 3 (0/3) |
| Tg2576 (APP<sup>+/-</sup>) | haPrP |  |  | 3 (0/3) | 5 (0/5) | 4 (0/4) |
| **<u>Controls</u>** |  |  |  |  |  |  |
| APP/PS1 (APP<sup>-/-</sup>) |  |  |  | 2 (2/0) | 11 (7/4) | 2 (0/2) |
| hAPP-J20 (APP<sup>-/-</sup>) |  |  | 2 (1/1) |  | 17 (9/8) | 1 (0/1) |

Transgenic mouse lines used in this study and the number of experimental animals in each group. Numbers in parentheses indicate the number of males and females in each mouse line/age group combination. The promoter used to drive APP expression for each transgenic line is indicated in the table. Control mice were APP<sup>-/-</sup> littermates of experimental mice.

### Two-photon serial imaging of methoxy-X04 labeled plaques

To label plaques, mice received an intraperitoneal (i.p.) injection of 3.3 mg/kg methoxy-X04 in 3.3% DMSO, 6.7% Kolliphor-EL (Millipore Sigma) in PBS. Twenty to twenty-four hours after injection, mice were perfused with 4% paraformaldehyde (PFA, 4ºC), then brains were dissected and post-fixed in 4% PFA at room temperature for 3-6 hours, followed by overnight at 4°C. Whole brain fluorescence imaging was performed as described in (Oh et al., 2014) with serial two-photon (STP) tomography (Ragan et al., 2012; TissueCyte 1000, TissueVision Inc. Somerville, MA), using 925 nm excitation, a 500 nm dichroic mirror, and a 447/60 bandpass emission filter on the blue channel. Serial block-face images were acquired at 0.35 μm/pixel lateral resolution with a 100 μm sectioning interval. We acquired 140 serial sections through each brain from cerebellum through olfactory bulb.

### Segmentation and registration

Automated image segmentation was performed as previously described (Kuan et al., 2015) with the following modifications. Candidate plaque areas were identified by performing adaptive thresholding on band-passed blue channel pixel strength in relation to the relative signal strength in green channel. This additional step was implemented because many artifacts with detectable blue signal tended to have relatively lower green signals than that of true plaques identified by expert annotation. Since artifacts were more prevalent around tissue borders and in ventricles, the candidate plaques are then further probabilistically filtered by a simple morphometric classifier which measures and tests the object shape elongation, spatial location/distance to tissue border, and its relative signal strength to tissue background autofluorescence in both green and red channels. Thirty-five of one hundred eleven image series were acquired with a lower photomultiplier tube (PMT) voltage (below 750 V) initially and were then processed with a higher sensitivity initial classifier.

Automated 3D registration was also performed as previously described (Kuan et al., 2015). Briefly, segmented fluorescence output is a full resolution mask that classifies each 0.35 μm × 0.35 μm pixel as either signal or background. An isotropic 3D summary of each brain is constructed by dividing each image series into 10 μm × 10 μm × 10 μm grid voxels. Total signal is computed for each voxel by summing the number of signal positive pixels in that voxel. Each image stack is registered in a multi-step process using both global affine and local deformable registration to the 3D Allen Mouse Brain Common Coordinate Framework, v3 (CCFv3). Plaque density for each structure in the reference atlas ontology was calculated by summing voxels from the same structure. We also used a standard feature labeling algorithm to obtain plaque counts within each structure. Adjacent and orthogonally adjacent voxels in the segmentation signal were grouped together as one plaque object. Due to the 100 μm z-sampling interval, our resolution limit for detecting separate plaques in the z axis was 100 μm.

### Quality Control

All image series were subjected to manual QC checks for completeness and uniformity of raw fluorescence images, minimum fluorescence intensity, and artifacts. Severe artifacts such as missing tissue or sections, poor orientation, edge cutoff, tessellation, and low signal strength caused image series failure. Brains that contained slices or other damage from the stereotaxic injection, dissection, and imaging/sectioning process were failed if the damage was in the right hemisphere. APP^−/−^ brains did not undergo additional QC checking beyond the raw image series. For APP^+/−^ brains, automatic segmentation results were checked for overall quality and false positive signals using a two-step process. First, every image series was manually scored by an expert annotator by overlaying segmentation results for 3-5 single coronal sections with raw fluorescent images from STP imaging. A qualitative score (from 1-7) was assigned to each image series based on the perceived overlap of the segmentation and raw image and absence of artifacts. Second, 3D gridded plaque images for every brain were loaded in ITK-SNAP (Yushkevich et al., 2006) and checked for obvious artifacts at tissue edges. Artifacts identified in 3D images were subsequently checked by overlaying the corresponding single sections. In some cases, (~25), the quality control process extended to identification and masking of areas of high intensity/high frequency artifacts and areas of signal dropout. The edge of the cerebral aqueduct, the cerebellum, and the medial border of the orbital cortex were particularly prone to bright tissue edge effects (**Figure 2f-k**), so these regions were manually checked in 1-5 coronal sections and the 3D grid file for every image series. In some cases, false positives in these regions could not be masked due to overlapping true positive signal and manual annotators made a judgment call for inclusion based on the overall quality of the image series. In general, APP^+/−^ segmentations were failed for false positives in rostral cortex and severe false positives in ventricles, but not for minor false positive signal in the ventricles, cerebellum and olfactory bulb.

**Table 2.** Abbreviations and full structure names for all annotated isocortical, hippocampal, cortical subplate, and olfactory areas in the Allen Mouse Brain Common Coordinate Framework (CCF) at the summary structure level.

| **Abbreviation** | **Full structure name** |
| --- | --- |
| **Isocortex** |  |
| FRP | Frontal pole, cerebral cortex |
| MOs | Secondary motor area |
| ACA<sub>d</sub> | Anterior cingulate area, dorsal part |
| ACA<sub>v</sub> | Anterior cingulate area, ventral part |
| PL | Prelimbic area |
| ILA | Infralimbic area |
| ORBI | Orbital area, lateral part |
| ORB<sub>m</sub> | Orbital area, medial part |
| ORB<sub>vl</sub> | Orbital area, ventrolateral part |
| Ald | Agranular insular area, dorsal part |
| Alv | Agranular insular area, ventral part |
| Alp | Agranular insular area, posterior part |
| GU | Gustatory areas |
| VISC | Visceral area |
| SSs | Supplemental somatosensory area |
| SSp-bfd | Primary somatosensory area, barrel field |
| SSp-tr | Primary somatosensory area, trunk |
| SSp-ll | Primary somatosensory area, lower limb |
| SSp-ul | Primary somatosensory area, upper limb |
| SSp-un | Primary somatosensory area, unassigned |
| SSp-n | Primary somatosensory area, nose |
| SSp-m | Primary somatosensory area, mouth |
| MOp | Primary motor area |
| VIS<sub>al</sub> | Anterolateral visual area |
| VIS<sub>l</sub> | Lateral visual area |
| VIS<sub>p</sub> | Primary visual area |
| VIS<sub>pl</sub> | Posterolateral visual area |
| VIS<sub>li</sub> | Laterointermediate area |
| VIS<sub>por</sub> | Postrhinal area |
| VIS<sub>rl</sub> | Rostrolateral visual area |
| VISa | Anterior area |
| VISam | Anteromedial visual area |
| VISpm | posteromedial visual area |
| RSPagl | Retrosplenial area, lateral agranular part |
| RSPd | Retrosplenial area, dorsal part |
| RSPv | Retrosplenial area, ventral part |
| AUDd | Dorsal auditory area |
| AUDp | Primary auditory area |
| AUDpo | Posterior auditory area |
| AUDv | Ventral auditory area |
| TEa | Temporal association areas |
| ECT | Ectorhinal area |
| **Hippocampal Formation** |  |
| CA1 | Field CA1 |
| CA2 | Field CA2 |
| CA3 | Field CA3 |
| DG | Dentate gyrus |
| FC | Fasciola cinerea |
| IG | Induseum griseum |
| ENTl | Entorhinal area, lateral part |
| ENTm | Entorhinal area, medial part, dorsal zone |
| PAR | Parasubiculum |
| POST | Postsubiculum |
| PRE | Presubiculum |
| SUB | Subiculum |
| ProS | Prosobiculum |
| HATA | Hippocampo-amygdalar transition area |
| APr | Area prostriata |
| **Olfactory Areas** |  |
| MOB | Main olfactory bulb |
| AOB | Accessory olfactory bulb |
| AON | Anterior olfactory nucleus |
| TT | Taenia tecta |
| DP | Dorsal peduncular area |
| PIR | Piriform area |
| NLOT | Nucleus of the lateral olfactory tract |
| COAa | Cortical amygdalar area, anterior part |
| COAp | Cortical amygdalar area, posterior part |
| PAA | Piriform-amygdalar area |
| TR | Postpiriform transition area |
| **Cortical Subplate** |  |
| CLA | Clastrum |
| EPd | Endopiriform nucleus, dorsal part |
| EPv | Endopiriform nucleus, ventral part |
| LA | Lateral amygdalar nucleus |
| BLA | Basolateral amygdalar nucleus |
| BMA | Basomedial amygdalar nucleus |
| PA | Posterior amygdalar nucleus |

### Antibody Characterization

**6E10** (Covance #B228658, RRID: AB_1977025, 1:1000) This mouse monoclonal IgG1 antibody is reactive to amino acid residues 1-16 of human β amyloid, specifically recognizing the epitope of amino acids 3-8 of the sequence (EFRHDS). This antibody has been previously shown to label amyloid plaques in the brains of humans (Patton et al., 2006) and all three AD transgenic mice used in this study (Y. Huang et al., 2015; Pozueta et al., 2013; Thakker et al., 2009). No staining was observed when the antibody was used to stain tissue from APP^−/−^ mice. ***Goat Anti-Mouse IgG1(Alexa Fluor^®^ 568).*** This goat polyclonal IgG1 antiserum (Invitrogen #1964384, RRID: AB_2535766, 1:1000) is reactive against mouse IgG1. No staining was observed in tissue that was left unexposed to 6E10 primary antibody but incubated with the IgG1 secondary, for mice of all genotypes.

### Immunohistochemistry

Coronal sections retrieved after two-photon serial imaging (100 μm thickness) were immunostained to assess Aβ load and spatial distribution. For antigen retrieval, sections were placed in 70% formic acid for 15 minutes, followed by a PBS rinse. All sections were then incubated in blocking solution (4% normal goat serum + 0.5% Triton X-100 in PBS) for 2 hours. After blocking, sections were stained with primary antibody overnight (6E10, Covance, Princeton, NJ, Lot #B2286581:1000). After three 2-hour washes in PBS + 0.1% Triton X-100, sections were then incubated in secondary antibody overnight (Alexa Fluor goat anti-mouse 568). After three additional 2-hour washes in PBS + 0.1% Triton X-100, sections were counterstained with DAPI (Invitrogen, Carlsbad, CA, Lot #1874814) and coverslipped with Fluoromount G medium (Southern Biotechnology, Birmingham, AL; catalog #J3017-XE67B). All slides were imaged on the VS120 multichannel epifluorescence microscope system (Olympus, Center Valley, PA) with a 10X objective. Selected ROIs were subsequently imaged on the confocal laser scanning system FV3000 (Olympus, Center Valley, PA).

### Image Quantification

To quantify the fraction of methoxy-X04 labeled plaques detected by the automated segmentation algorithm following STP imaging, we manually counted the number of plaques in 1.4 mm x 1.4 mm ROIs in raw STP coronal images and their associated segmentation mask files (n=15 ROIs total, 3 ROIs per experiment in 5 experiments). All ROIs were drawn in the isocortex and hippocampus in 13-month-old or 19-month-old brains (n=3 APP/PS1, n=1 Tg2576, n=1 hAPP-J20). We also quantified the fraction of antibody-labeled plaque area detected by STP imaging and automated segmentation of methoxy-X04 labeling. We drew ROIs corresponding to major brain divisions on two sections from a single brain for each mouse line, then applied a threshold to the 6E10 antibody labeling using Fiji (**Figure 4c**). The area in the antibody-labeled image and the corresponding segmentation of the STP image were measured for each ROI, and the fraction detected was quantified as area in segmentation of the STP image / area in thresholded IHC image. To quantify the ratio of dense core to diffuse plaques, we drew polygons around the edge of the methoxy-X04 labeling and the Aβ antibody fluorescence in maximum intensity projections of confocal images (n=6 plaques per brain, 3 cortical and 3 hippocampal) and measured the area inside each polygon using Fiji (Schindelin et al., 2012). We used sections from one 13-month-old APP/PS1 mouse, one 13-month-old hAPP-J20 mouse, and one 19-month-old Tg2576 mouse.

### Statistics

Previous studies have reported that plaque density is not normally distributed (Liu et al., 2017). We performed the Shapiro-Wilk test for normality on the 13-month-old APP/PS1 group since it was the only group of mouse line and age having at least 30 samples (Razali & Wah, 2011). The test rejects the normality of the brain-wide plaque density distribution with a p-value of 0.001. Therefore, unless otherwise specified in the text, we used a Kruskal-Wallis one-way ANOVA with a significance level of 0.05 and Wilcoxon rank-sum test for post hoc comparisons for all hypothesis testing herein and we report plaque densities as median ± interquartile range (IQR).

## 3. Results

We measured the brain-wide distribution of plaque pathology in three AD mouse models that express mutant forms of APP using a high-throughput, high resolution imaging and analysis pipeline (**Figure 1a**). First, plaques were labeled *in vivo* with i.p. injection of methoxy-X04 one day before transcardial perfusion (Klunk et al., 2002). Methoxy-X04 crosses the blood brain barrier, labeling amyloid brain-wide and producing bright fluorescence that is natively detectable; properties critical for use in our STP pipeline (Amato, Pan, Schwartz, & Ragan, 2016). Methoxy-X04 primarily fluoresces in the blue channel, but, at least given our acquisition parameters, some signal was also detected in green and red channels. Some of this signal appears to be autofluorescence, as low, but still detectable, plaque signal was also observed in APP+/- mice that did not receive a methoxy-X04 injection as has been previously reported (Diez, Koistinaho, Kahn, Games, & Hökfelt, 2000; Dowson, 1981; Kwan, Duff, Gouras, & Webb, 2009; D R Thal, Ghebremedhin, Haass, & Schultz, 2002; Zipfel et al., 2003, **Figure 1b**). To establish a time course for plaque deposition in each transgenic mouse line, methoxy-X04 injections were performed in multiple age groups between 5 and 19 months (**Table 1**). Second, following plaque labeling, brains were imaged with serial two-photon tomography (STP) at high x,y resolution (0.35 × 0.35 μm) every 100 μm throughout the entire rostral-caudal extent of the brain in coronal planes (**Figure 1b**). Next, each whole brain image series was processed using the informatics pipeline adapted from (Kuan et al., 2015). This step consists of two parts; signal detection (segmentation) and registration. We developed a custom-built signal detection algorithm to automatically segment the methoxy-X04 labeled plaques in each serial section (**Figure 2,** see methods for details). Segmented image stacks were deformably registered to the 3D Allen Mouse Brain Common Coordinate Framework, v3 (CCFv3) and resampled to 10 μm voxel resolution (“3D grid files”). Finally, we quantified plaque density and/or plaque number from the methoxy-X04 signal and automated segmentation for every region annotated in the Allen CCFv3 reference atlas across the entire brain.

**Figure 1.**
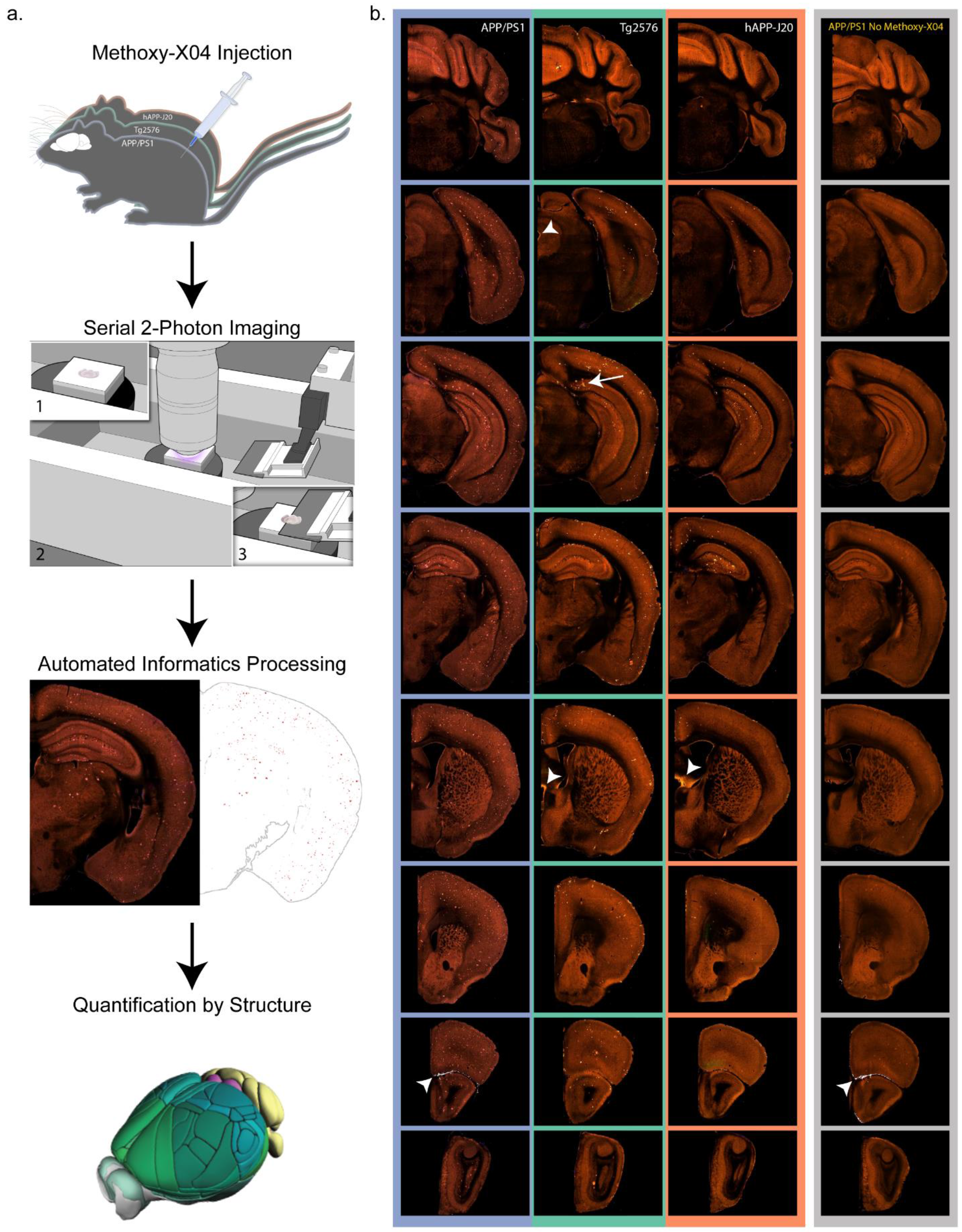
Pipeline workflow for labeling and quantification of brain-wide plaque distribution. **(a)** Mice were injected with methoxy-X04 to label plaques *in vivo* 24 hours before perfusion. Brains were imaged with serial 2-photon tomography which involves embedding the brain in agar (1), acquisition of 2-photon images in the coronal plane (2), and sectioning with an integrated vibratome (3). Images were processed in an automated informatics pipeline that included automated detection of plaques from section images and deformable 3D registration to the Allen CCFv3. Plaque density and count were then quantified for all structures. **(b)** Selected images from coronal STP image series spanning the brain from cerebellum to olfactory bulb are shown for a 19-month-old mouse with methoxy-X04 labeling from each of the three APP mouse lines used in this study: APP/PS1, Tg2576, and hAPP-J20. Images in the furthest right column show similar sections from a 6-month-old APP/PS1^+/−^ mouse that was not injected with methoxy-X04. Some lower intensity plaque fluorescence is visible without methoxy-X04 labeling. Arrow indicates methoxy-X04 labeled plaques in the subiculum of Tg2576 mice. Arrowheads show examples of bright tissue edges that can cause false positives in automated segmentation.

**Figure 2.**
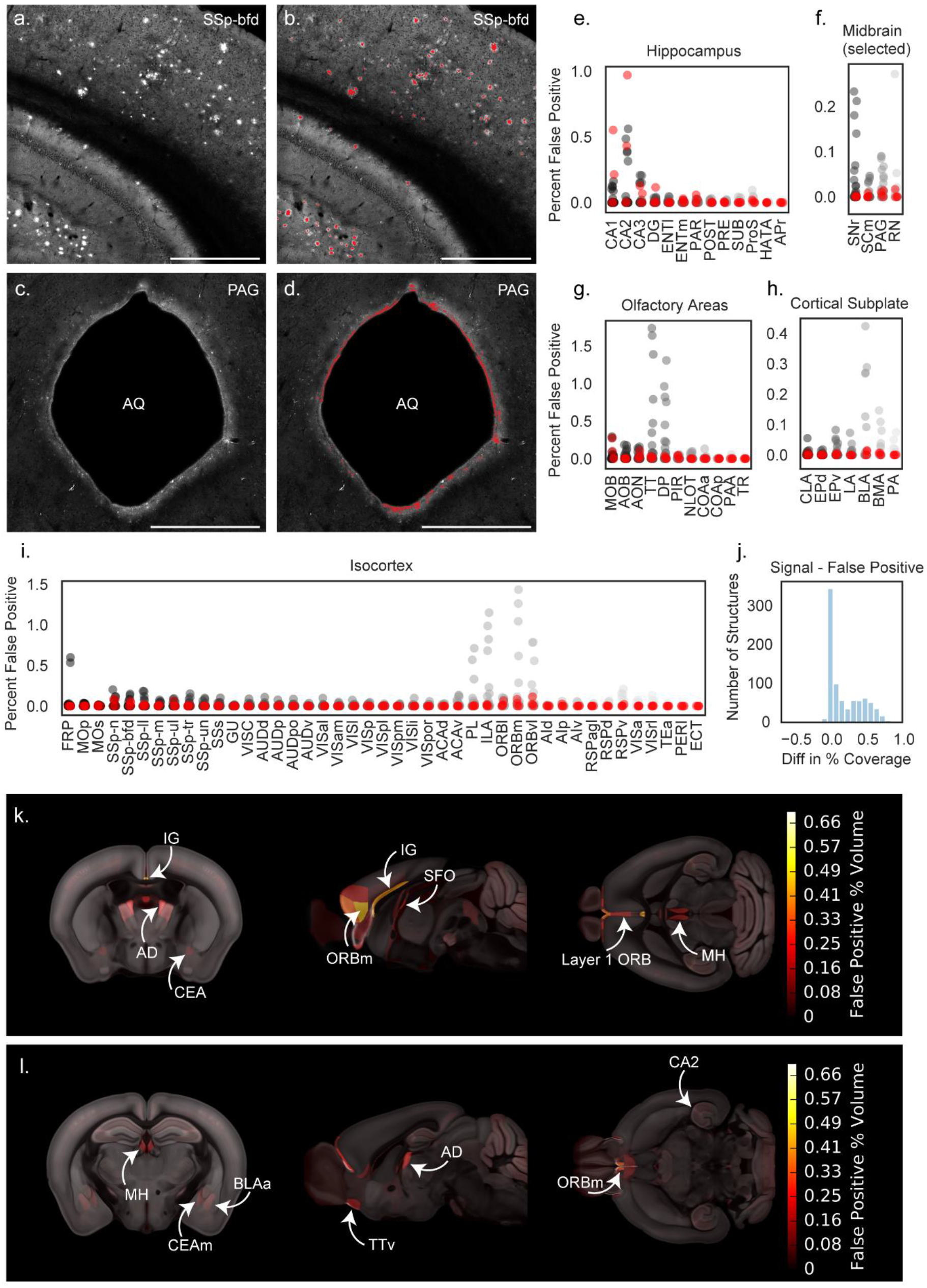
Automated segmentation of plaque fluorescence. **(a)** An example ROI from a single section image in somatosensory cortex, barrel field area (SSp-bfd) showing methoxy-X04 fluorescence. A portion of the hippocampus is also visible in the bottom left corner of the image. **(b)** Segmented plaque signal (red) overlaid on the image section shown in (a). **(c)** An ROI from a different section in the same brain showing the cerebral aqueduct (AQ) and surrounding periaqueductal gray (PAG). **(d)** Segmented plaque signal for the image section shown in c (red) showing false positive signal along the bright tissue edge at the border of the cerebral aqueduct. Similar signal was observed in APP^−/−^ mice. Scale = 1 mm. **(e-i)** Quantification of false positives as percent of structure volume for 35 image series collected from APP^−/−^ mice without plaques to identify regions prone to segmentation artifacts. False positives are plotted for summary structures in hippocampus **(e)**, selected midbrain regions including PAG **(f)**, olfactory areas **(g)**, cortical subplate **(h)**, and isocortex **(i)**. Red points indicate APP^−/−^ mice that received a methoxy-X04 injection one day before perfusion. Abbreviations for summary structures in isocortex, hippocampus, olfactory areas, and cortical subplate can be found in **Table 2**. Abbreviations in (f) SNr: Substantia nigra, reticular part; SCm: Superior colliculus, motor related; PAG: Periaqueductal gray; RN: Red nucleus. **(j)** Histogram showing the difference between signal and false positive levels (% coverage in APP^+/−^ mice - % coverage in APP^−/−^ mice) for every structure in the Allen CCFv3 reference atlas. **(k)** False positive heatmap showing regions with the highest segmentation artifacts. Abbreviations: BLAa: Basolateral amygdalar nucleus, anterior part; IG: Induseum griseum; AD: Anterodorsal nucleus of the thalamus; CEA: Central amygdalar nucleus; ORBm: Medial orbital cortex; SFO: Subfornical organ; MH: Medial habenula. (l) False positive heatmap for a different anatomical location. Abbreviations: CEAm: Central amygdalar nucleus, medial part; TTv: Taenia tecta, ventral part; CA2: Hippocampal field CA2.

### 3.1 Evaluation of automated plaque segmentation

The performance of the segmentation algorithm was analyzed in two ways. First, automatic segmentation results were manually scored for overall quality by an expert annotator (see methods for details). Second, we compared the number of methoxy-X04 labeled plaques detected by the automated segmentation algorithm with the number of plaques manually identified on the raw images for a subset of experiments and regions of interest (ROIs). On average, the automated segmentation algorithm produced slightly higher plaque counts compared to manual counts (115+/−33%). This discrepancy is likely due in part to the observation that some brain areas were particularly prone to false positives in the segmentation, most often caused by bright tissue edges. These artifacts were commonly seen in the cerebellum, around the edges of ventricles, particularly the cerebral aqueduct (**Figure 2d)**, in the rostral cortex, particularly layer 1 of orbital cortex, and in the glomerular layer of the olfactory bulb where lipofuscin deposits could not be distinguished from methoxy-X04-labeled plaques. The segmentation and 3D grid files were manually checked for every brain, with particular attention paid to these artifact-prone areas.

Since some regions of the brain were prone to segmentation artifacts, confidence in automatically-generated quantification of plaque densities is lower for some regions than others. To determine which regions were the most problematic (and therefore had the lowest confidence), we subjected a set of 35 STP image series from wild type, non-transgenic APP^−/−^ littermates with and without methoxy-X04 injections, to the same analysis pipeline as our APP^+/−^ brains, *except for the segmentation QC*, so that we could directly measure the occurrence of artifacts contributing to false positives in our data. Following the application of the plaque segmentation algorithm to these brains, we quantified the percentage of voxels containing false positive signal for structures in the isocortex, hippocampus, midbrain, olfactory areas, and cortical subplate (**Figure 2e-i).** To estimate the magnitude of false positive signals across the whole brain, we calculated the relative error per structure by subtracting the mean signal in the control dataset from the mean signal in the full plaque dataset for every structure annotated in the Allen CCFv3 reference atlas (839 structures). The mean false positive signal for every structure is plotted in **Figure 2j** and included in **Appendix 1.** Fifty-two percent of structures had an error between −0.1 and 0.1 indicating that less than 0.1% of their signal was affected by false positives. There was a second, smaller peak in the histogram around 0.5%, composed of structures that had some false positive signal. Sixteen percent of structures had more than 0.5% difference between their plaque density and their false positive density, and no structure had more than a 0.9% difference (**Figure 2j**). Sixteen percent of the 839 structures had small negative values for the difference between signal and false positive percent coverage, indicating higher false positives than true plaque signal. However, it is important to note that the control dataset did not undergo the same segmentation QC process as the plaque dataset, so the reported false positive values are an upper bound on the error. To make it easier to visualize the anatomical locations of regions with high false positive signal, we generated a false positive heat map showing the spatial distribution of regions with high segmentation background (**Figure 2k,l**). Most of these regions are located along the midline or near ventricles and are subject to false positives from bright tissue edges (arrowheads in **Figure 1**).

### 3.2 Metrics for quantification of plaque load using methoxy-X04 and comparison to Aβ antibody labeling

We explored several features for quantifying plaque pathology. Most frequently, we report plaque density per structure, defined as the percent of each structure’s total volume that contains segmented plaque signal. We also calculated plaque counts based on the segmentation results. To accomplish this, we computationally identified voxels with segmented plaque that were touching and/or diagonally adjacent to each other in the 3D grid images (**Figure 3**). Our 100 μm z-sampling interval caused plaques to have an elongated appearance along the anterior-posterior axis (**Figure 3c,d**). However, we also identified individual plaques that extended for several millimeters across serial sections, much further than our sampling interval would explain. This was particularly apparent along the midline where many plaques appeared to follow the path of the vasculature. This was more obvious in hAPP-J20 and Tg2576 mice (**Figure 3e, f**) than in APP/PS1 mice (**Figure 3d**), but was observed in all three APP mouse lines. Because we could not easily differentiate between these extended length vascular-associated plaques and the non-vascular plaques based on the automated counting results, in most of our subsequent analyses we report plaque density to describe pathological load. However, the number of plaques per cubic mm are reported in **Figure 5a** and included in **Appendix 2**, and it should be noted that these values were not corrected for the presence of vascular-associated plaques.

**Figure 3.**
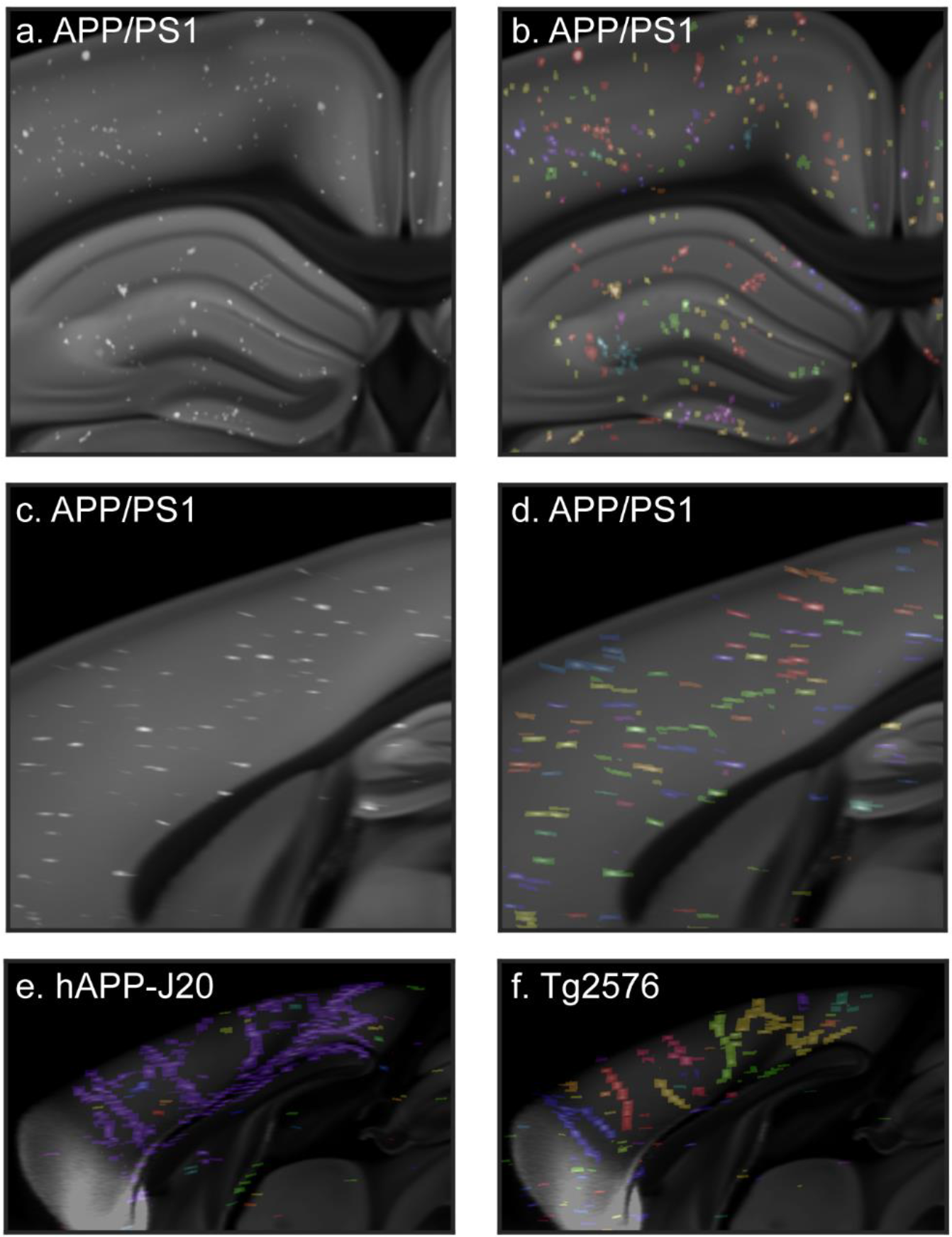
Automatic plaque counts reveal large, vascular-associated amyloid deposits. Portion of a single coronal **(a,b)** and sagittal **(c,d)** image from the CCFv3 template with gridded, automatically detected methoxy-X04 labeled plaques from a 19-month-old APP/PS1 mouse brain overlaid in white. In (**b**) and **(d)**, the same images are shown with automatically counted plaques in labeled with different colors. Plaques in the same area that are the same color had diagonally adjacent voxels in at least one image section and were counted as one single object. The sagittal view in (d) shows that plaques are elongated on the z-axis. **(e)** Sagittal image from the CCFv3 template with gridded plaques from a 19-month-old hAPP-J20 mouse with automatic plaque count results overlaid. Nearly all of the plaques along the midline in this brain were counted as a single plaque (blue). **(f)** Sagittal CCFv3 template image with gridded plaques and counts overlaid for a 19-month-old Tg2576 mouse reveals large, vascular-associated deposits.

Methoxy-X04, a congo red derivative, was reported to label dense-core amyloid but not diffuse plaques (Condello, Schain, & Grutzendler, 2011). Thus, we wanted to characterize the relative fraction of *total* plaque detectable with methoxy-X04 compared to labeling sections with an Aβ antibody. We retrieved individual sections from a subset of experiments following methoxy-X04 labeling and STP imaging, and immunostained them with the 6E10 anti-Aβ antibody (**Figure 4a,b**). We quantified the area covered by plaques in thresholded, antibody-labeled sections compared with the automated segmentation of the same sections **(Figure 4c**, *e.g.,* compare red to white masks). In all sections, the overall plaque density in the antibody-labeled sections was higher than the automated segmentation of methoxy-X04 fluorescence, but the densities for the two methods of plaque labeling differed between mouse lines (APP/PS1: 10.4±7.8% IHC, 1.2±0.80% segmentation (13-months); Tg2576: 2.8±2.5% IHC, 0.44±0.37% segmentation (19-months); hAPP-J20: 7.2±7.2% IHC, 0.08±0.07% segmentation (13-months)). To more directly measure our detection level, we calculated the fraction of Aβ antibody-labeled area that was detected by automatic segmentation of methoxy-X04 fluorescence in the same section for all ROIs in each image. Surprisingly, we found a significant effect of mouse line on the fraction of antibody-labeled plaque area detected (**Figure 4f,** p=0.04, one-way ANOVA). In APP/PS1 and Tg2576 mouse lines, the fraction detected by segmentation was similar (0.15±0.10 for APP/PS1, 0.15±0.11 for Tg2576), but a lower proportion of antibody labeled plaques were detected in hAPP-J20 (0.03±0.03, p=0.04 compared to APP/PS1, Tukey’s post hoc comparison). This line-specific difference may be due to the significantly lower ratio of dense core to diffuse plaque area that we measured in hAPP-J20 mice compared to the other two lines (APP/PS1: 0.09±0.04, Tg2576: 0.1±0.03, hAPP-J20: 0.03±0.02, p=0.001 one-way ANOVA, **Figure 4g**). We did not observe any antibody-labeled plaques in regions without methoxy-X04 labeled plaques, and critically, plaque densities measured by both methods are very highly and significantly correlated for all regions tested in all three mouse lines (Pearson’s r=0.95, p<0.0001, **Figure 4e**), supporting the use of the methoxy-X04 label in our systematic pipeline approach to describe brain-wide distribution of plaques. Thus, we find that methoxy-X04 does underestimate total, or absolute, plaque density (including diffuse plaques) by ~7- to 10-fold for APP/PS1 and Tg2576 mice, and ~30-fold for hAPP-J20 mice. Due to this difference in the fraction of plaques labeled with methoxy-X04, plus known differences in the rate of plaque accumulation in each mouse line (Jankowsky & Zheng, 2017), we focused our analysis on *patterns* of plaque deposition rather than comparing absolute levels between lines. Where we do compare plaque levels (**Section 3.3**), it is important to note that these reported values refer only to dense-core, not diffuse, plaques.

**Figure 4.**
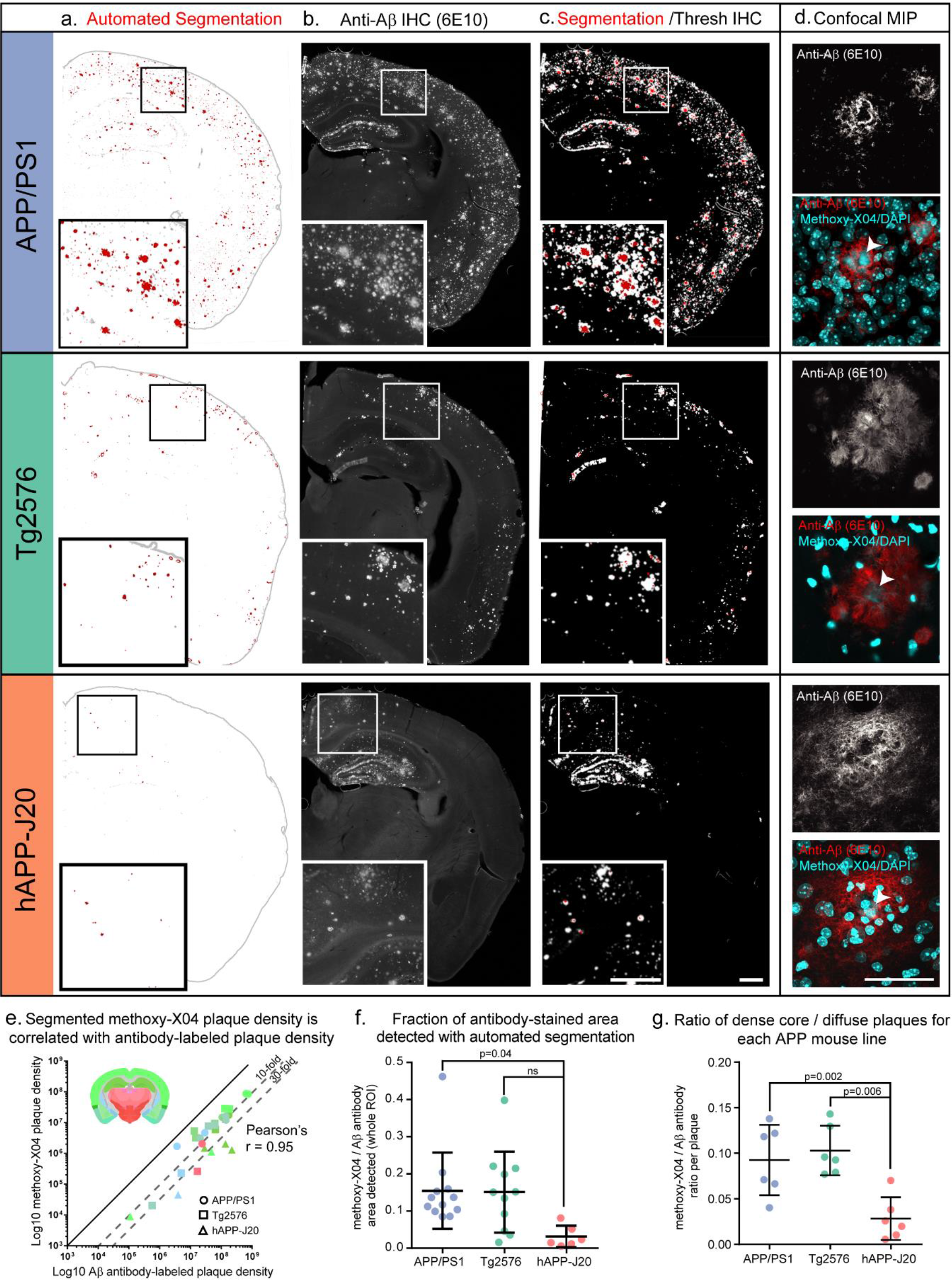
Methoxy-X04 accurately labels the location of plaques but underestimates plaque density. **(a-c)** Automated segmentation results **(a)**, IHC for Aβ (6E10 antibody) **(b)**, and an overlay of the automated segmentation and thresholded IHC image from the same section **(c)** are shown for one section from each mouse line. APP/PS1 and hAPP-J20 brains are from 13-month-old mice. The Tg2576 brain is from a 19-month-old mouse. In the segmentation images in (a), red indicates segmented plaques, and gray the putative plaque signal that was removed by the adaptive filtering step in the segmentation algorithm. Automatically detected tissue edges are also colored gray. Image alignment in the overlay images **(c)** is imperfect due to tissue deformation during antibody labeling. **(d)** maximum intensity projection of a confocal image stack through the center of one plaque for each mouse line. Upper grayscale image shows 6E10 labeling; bottom color image shows 6E10 labeling overlaid with methoxy-X04 and DAPI fluorescence (both in cyan). Arrowheads indicate the methoxy-X04 positive core of each plaque. Scale = 500 μm (a-c), 50 μm (d). **(e)** Aβ antibody-labeled plaque density and segmented methoxy-X04-labeled plaque density plotted for each ROI measured (log scale). Points are colored by the Allen CCFv3 reference atlas region in which each ROI was drawn (isocortex, hippocampal formation, cortical subplate, olfactory areas, striatum, and thalamus). Inset shows a coronal section from the Allen CCFv3 Reference Atlas with region colors. The shape of each point indicates which mouse line it belongs to: circles for APP/PS1, squares for Tg2576, and triangles for hAPP-J20. Methoxy-X04 density was lower than Aβ antibody-labeled density for every point, but the two measurements were highly correlated (Pearson’s r = 0.95, p<0.0001). **(f)** The fraction of Aβ antibody-labeled plaque density that was detected by automated segmentation was significantly lower for hAPP-J20 mice compared to APP/PS1 mice. **(g)** For individual plaques, the ratio of methoxy-X04 labeling to Aβ antibody labeling is plotted for each mouse line. hAPP-J20 mice had significantly lower fractions of methoxy-X04/Aβ antibody labeling.

### 3.3 Comparison of methoxy-X04-labeled plaque levels between three APP mouse models

We measured methoxy-X04 labeled plaque density and count for each mouse line across multiple ages in the whole brain (**Figure 5a,b**). Of the three lines, APP/PS1 mice showed the most aggressive rates of dense-core plaque accumulation, as expected from previous reports (Garcia-Alloza et al., 2006; Jankowsky et al., 2004). Whole-brain dense core-plaque densities were significantly higher in APP/PS1 mice compared to the other two mouse lines in every age group, and whole brain plaque counts were higher in APP/PS1 mice compared to the other two lines for every age except 5 months. At 5 months-old, plaques were readily observable in APP/PS1 mice. Occasionally, but still quite infrequently, plaques were detected in 5 and 7-month-old hAPP-J20 mice. At 13 months, hAPP-J20 mice had significantly higher plaque density than Tg2576 mice (**Figure 5b**, p=0.04). Very sporadic plaques were observed in Tg2576 mice at 13 months, and they then accumulated more plaques rapidly between 13 and 19 months.

**Figure 5.**
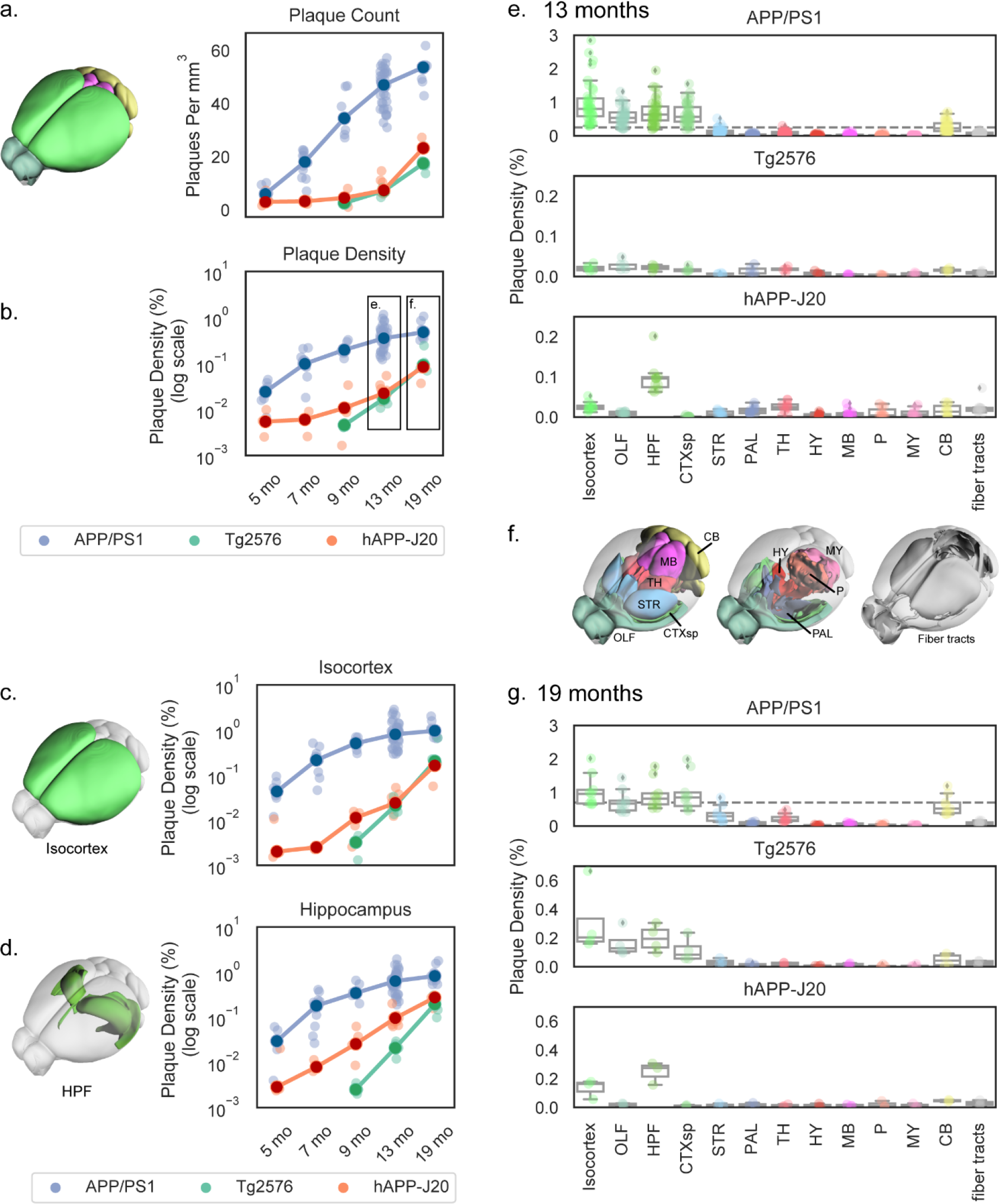
Automated quantification of methoxy-X04-labeled plaque density and count across whole brain and major divisions for three APP mouse lines at various ages. **(a,b)** Plaque density measured by automated segmentation and registration plotted as count (plaques per mm^3^; **a**) or density (% of structure volume; **b-g**). Plaque density is plotted by age for the whole brain **(a,b)**, isocortex **(c)**, and hippocampal formation **(d)** in each of the three APP lines characterized. Dark circles indicate the median values and light points individual animals. Lines connect points at the median. Plaque density at 13 months **(e)** and 19 months **(g)** divided by major brain division is plotted separately for each mouse line. Box plots show the median and IQR, with whiskers extending to 1.5 times the IQR. Outliers are plotted as individual points beyond the whiskers. Dashed vertical line in the APP/PS1 plots in (e) and (g) show the maximum value from the hAPP-J20 and Tg2576 mouse line plots below at the same age. Plots in (e) and (g) are colored by major brain division, and the cartoons in (c), (d), and (f) show the anatomical locations of the major brain divisions in the Allen CCFv3 reference atlas. Abbreviations: OLF: Olfactory areas, HPF: Hippocampal formation, CTXsp: Cortical subplate, STR: Striatum, PAL: Pallidum, TH: Thalamus, HY: Hypothalamus, MB: Midbrain, P: Pons, MY: Medulla, CB: Cerbellum.

We also measured methoxy-X04-labeled plaque densities within each of the twelve major brain divisions, and fiber tracts, annotated in the Allen CCFv3 reference atlas. In **Figure 5c,d**, we show plaque density across all age groups for the isocortex (**Figure 5c**) and hippocampal formation (**Figure 5d**), As seen at the whole brain level, methoxy-X04 labeled dense-core plaque densities in these two major regions were also higher in APP/PS1 mice compared to Tg2576 or hAPP-J20 mice at every age. Tg2576 and hAPP-J20 mice did not differ in isocortex plaque density, but hAPP-J20 mice had significantly higher plaque density in the hippocampal formation compared to Tg2576 mice at 9 and 13-months-old (p=0.02, p=0.04, **Figure 5d**).

### 3.4 Temporal and spatial progression patterns of methoxy-X04 labeled plaques in three APP mouse models

Plaque densities across all the major brain divisions are shown for all three mouse lines in 13- and 19- month-old brains in **Figure 5e-g**. APP/PS1 mice had the highest plaque density in the isocortex but also had dense plaques in olfactory areas, hippocampal formation, cortical subplate, and cerebellum at both 13- and 19-months-old. At 13 months, APP/PS1 mice were also beginning to show some plaque accumulation in the thalamus and striatum although the plaque density in these two structures was not significantly higher than the median for the whole brain at either 13 or 19 months. Tg2576 mice had only very sporadic plaques at 13 months, but by 19 months-old their plaque distribution across major regions looked similar to APP/PS1 mice; densest plaques in isocortex, followed by olfactory regions, hippocampal formation, cortical subplate, and cerebellum. However, even though the plaque distribution was broadly similar, Tg2576 still had much lower plaque densities than APP/PS1 mice (note the different scales for the APP/PS1 mouse line plot compared with Tg2576 and hAPP-J20 in **Figures 5e** and **5g**). Although there was a significant effect of structure on plaque density in major brain divisions in Tg2576 mice (p=10^-5^ at 13 months, p=0.001 at 19 months), no individual brain division had a significantly higher plaque density than the median in post hoc testing. Notably, hAPP-J20 mice had a different brain-wide plaque distribution from the other two lines. Plaque levels in the isocortex and hippocampal formation were significantly higher than the whole brain median at 13 months, and plaque density in the cortical subplate was significantly lower than the median at the same time point (**Figure 5e**). By 19 months, hAPP-J20 mice still had more plaques in the hippocampal formation than any other brain division and there was a significant effect of structure on plaque density (p=0.02), but none of the plaque densities in individual brain divisions were significantly different from the median at 19-months.

To examine the regional distribution of methoxy-X04 labeled plaques at a finer scale, we quantified densities for each of 316 subdivisions of the 12 major brain regions (aka, “summary structures”) in the Allen CCFv3. Plaque densities are plotted in **Figure 6** for summary structures in the 4 major brain divisions with the most plaques (isocortex, hippocampal formation, olfactory areas, and cortical subplate). Very small summary structures are not shown here (*i.e.,* total volume < 0.5% of its major division), but the plaque densities for all structures in all mouse lines and age groups are reported in **Appendix 2**.

**Figure 6.**
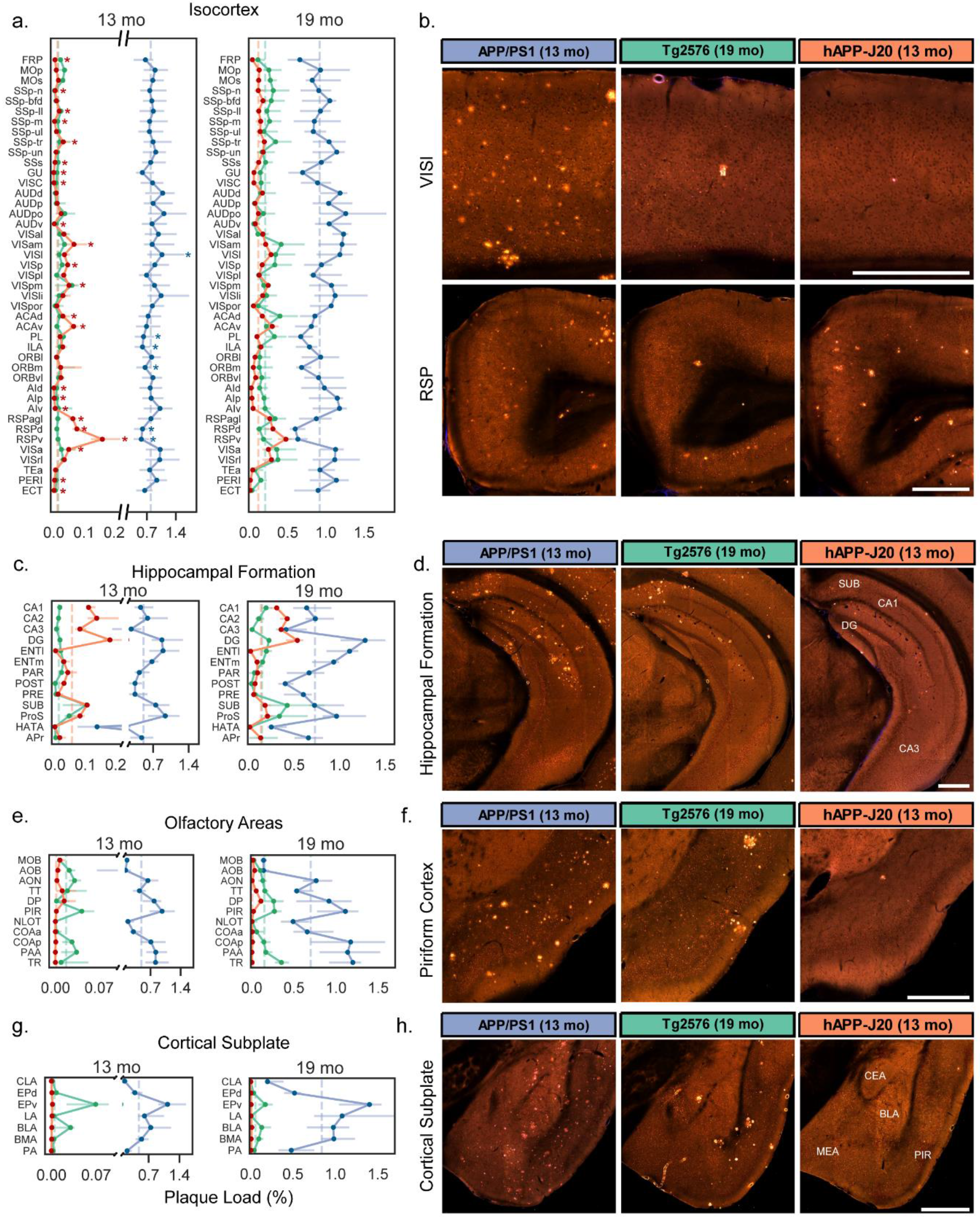
Quantification of plaque density for selected summary structures in the isocortex, hippocampus, olfactory areas, and cortical subplate. Plaque density (percent of structure volume) for summary structures in **(a)** isocortex, **(c)** hippocampal formation, **(e)** olfactory areas, and **(g)** cortical subplate in 13-month-old (left) and 19-month-old (right) mice. Darker lines connect points at the median and lighter error bars indicate the interquartile range. Dashed vertical lines indicate the median value for each mouse line for all graphed structures in each plot. Plots for plaque density at 13 months have split y-axes to allow all three datasets to be plotted on one graph. Asterisks in (a) indicate structures that were significantly higher or lower than the median plaque density in isocortex for APP/PS1 and hAPP-J20 mice. **(b)** Representative images showing plaque deposition patterns in each of the three APP mouse lines for two isocortex structures: lateral visual cortex (VISl, top) and retrosplenial cortex (RSP, bottom). **(d)** Images showing plaque deposition patterns in the hippocampus proper for the three mouse lines. The location of subiculum (SUB), CA1, CA3, and dentate gyrus (DG) are indicated on the hAPP-J20 image. **(f)** Images showing plaque deposition patterns in the piriform (olfactory) cortex for the three mouse lines. The hAPP-J20 image shows one of only a few plaques observed in this region in hAPP-J20 mice. **(h)** Images showing plaque deposition patterns in different amygdalar structures in the cortical subplate for the three mouse lines. The approximate location of the central amygdala (CEA), medial amygdala (MEA), basolateral amygdala (BLA), and piriform cortex (PIR) is indicated on the hAPP-J20 image. All abbreviations used in the graphs can be found in **Table 2**. Scale = 500 μm.

The isocortex is parcellated into 43 regions (or, summary structures) in the Allen CCFv3. We observed widespread distribution of plaques across these regions in APP/PS1 mice at 13 and 19 months (**Figure 6a)**. However, there was still some regional specificity within the cortex for this APP line. Plaque densities in the lateral visual area (VISl) were significantly higher than the median plaque density across the entire isocortex at 13 months (**Figure 6a,b**). Prefrontal areas prelimbic (PL), infralimbic (ILA), and medial orbital cortex (ORBm), had lower plaque density compared to the whole isocortex median (**Figure 6a**). hAPP-J20 mice showed a very striking and specific regional distribution of plaque in the isocortex at both 13 and 19 months, but we only performed statistical testing at 13 months since the isocortex was not identified as significant with post-hoc testing across all major divisions at 19 months. Plaque density in the isocortex of 13-month-old hAPP-J20 mice was extremely variable, with 23 summary structures having median plaque densities that were significantly higher or lower than the median for the whole structure (**Figure 6a**).

Specifically, we identified levels that were 5-13 times higher than the median in three subdivisions of retrosplenial cortex (RSPagl, RSPd, RSPv) at 13 months and 2-4 times higher than the median at 19 months, an observation consistent with previous reports (Harris et al, 2010). Interestingly, plaque density in RSPd and RSPv was significantly *lower* than the isocortex median for APP/PS1 mice (**Figure 6a,b**). In addition to RSP subdivisions, hAPP-J20 mice had high plaque density in ventral anterior cingulate cortex (ACAv), parietal association cortex (anterior visual area, VISa, anteromedial visual area, VISam, and posteromedial visual area, VISpm), and surprisingly, primary visual cortex (VISp). Tg2576 mice also appeared to have heavier plaque accumulation in dorsal anterior cingulate (ACAd) and parietal (VISam, VISa, and VISrl) cortex than in other cortical areas at 19 months (**Figure 6a**) but there was no significant effect of structure on plaque density in isocortex for this mouse line at any age.

The hippocampal formation (HPF) contains 14 summary structures (**Figure 6c**), including the hippocampus itself (CA1, CA3, DG, SUB) as well as associated regions (e.g., entorhinal cortex, ENT). Within the HPF, structure had a significant effect on plaque density for every mouse line/age group combination. In the hippocampus proper, all three lines had higher plaque density in the DG than in CA1, CA2, or CA3 (**Figure 6c,d**). APP/PS1 and Tg2576 mice also had high plaque density in lateral and medial entorhinal cortex (ENTl and ENTm). hAPP-J20 mice had plaques in ENTm but plaques in ENTl were rarely observed. All three mouse lines had relatively heavy plaque accumulation in the subiculum (SUB) and prosubiculum (ProS), but the Tg2576 mouse line had a strong preferential accumulation of plaques specifically in the dorsal subiculum (**Figure 6d,** white arrow in **Figure 1**). Notably, every single individual Tg2576 mouse that we imaged had plaques in the dorsal subiculum, even at 9 months-old.

Two other brain divisions with differences in plaque density between mouse lines were the olfactory areas which had dense plaques in many regions for APP/PS1 and Tg2576 mice, but virtually no plaques in hAPP-J20 mice (**Figure 1, Figure 6e,f**) and the cortical subplate which contains several amygdalar structures, where plaques were similarly high in APP/PS1 and Tg2576 but low or absent in hAPP-J20 mice (**Figure 6g,h**). Importantly, we also did not observe Aβ antibody-labeled plaques in olfactory regions or the amygdala in hAPP-J20 mice. Also of note, both APP/PS1 and Tg2576 mice had plaques in the cerebellum, whereas we did not identify any in hAPP-J20 mice (**Figure 7**). APP/PS1 mice developed cerebellar plaques as early as 4 months-old, but plaques were not observed in the cerebellum of Tg2576 mice until 19 months-old, and then many were associated with vasculature.

**Figure 7.**
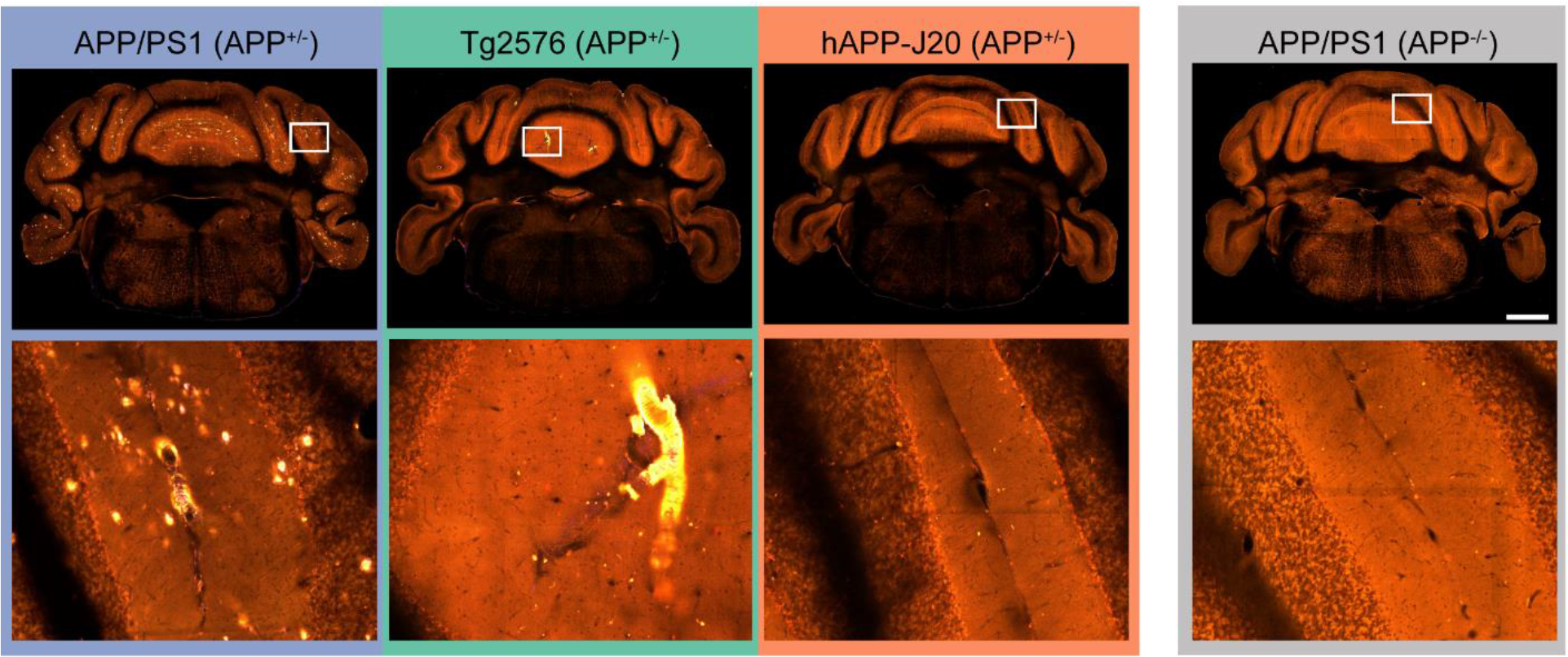
APP/PS1 and Tg2576 mice have plaques in the cerebellum but hAPP-J20 mice do not. Single STP image planes in the cerebellum (top row) and higher magnification views of the region indicated with the white box (bottom row). All images are from 19-month-old mouse brains. Scale = 1 mm.

We also explored the relationship between regional plaque distribution patterns in the isocortex with the structural connectivity-based isocortical modules found from a network analysis of modeled connectivity weights in wild type C57BL/6J mice (Harris et al., 2018, Knox et al., 2018). These modules are comprised of cortical regions that are more strongly connected to each other than to regions in other modules. The medial and prefrontal modules contain most of the regions reported to be part of the rodent DMN (Lu et al., 2012; Sforazzini, Schwarz, Galbusera, Bifone, & Gozzi, 2014; Stafford et al., 2014; Zerbi, Grandjean, Rudin, & Wenderoth, 2015); including RSP, ACA, posterior parietal (PTLp aka VISa and VISrl), ORB, and PL. The plaque density in each of the six cortical modules for each mouse line compared to the expected density if plaques were evenly distributed across the entire cortex is plotted in **Figure 8a**. hAPP-J20 mice had three times as many plaques in the medial module as would be expected with an even distribution and one and one-half times the expected density in the prefrontal module, indicating a distribution that overlaps with the rodent DMN. The relative plaque density in the medial module for hAPP-J20 mice was significantly higher than any other module (p<0.0001, 2-way ANOVA with Sidak’s multiple comparisons test), and relative plaque density in the prefrontal module was significantly higher than all other modules except visual (p<0.0001 for all other modules, p=0.5 for visual, 2-way ANOVA with Sidak’s multiple comparisons test). hAPP-J20 mice also had lower plaque density in the anterolateral, somatomotor, and temporal modules compared to all other modules, but these three modules did not differ from each other. Tg2576 mice had significantly fewer plaques in the temporal module compared to the prefrontal and visual modules (p=0.02, p=0.003, 2-way ANOVA with Sidak’s multiple comparisons test) and APP/PS1 mice had significantly fewer plaques in the medial compared to the visual module (p=0.003, 2-way ANOVA with Sidak’s multiple comparisons test). To visualize these differences in plaque distribution across modules, we created images of the Allen CCFv3 reference atlas structures showing the positions of the modules with a colormap corresponding to relative plaque levels for each of the three mouse lines (**Figure 8b**).

**Figure 8.**
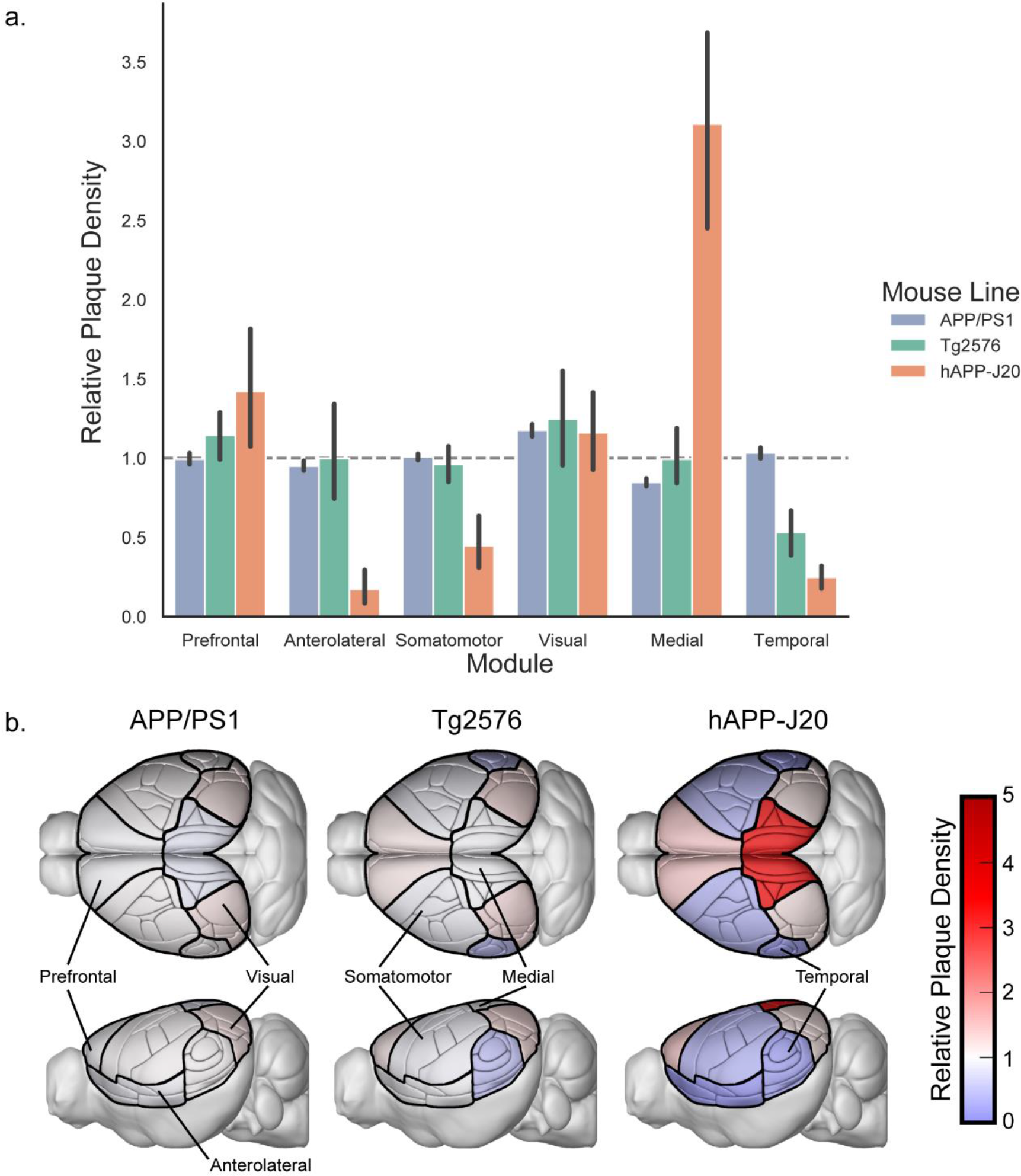
Plaques are unequally distributed across structural modules in the isocortex. **(a).** Plaque density in the isocortex modules from (Harris et al., 2018) relative to the density that would be expected with even plaque distribution across all cortical regions (dashed gray line). Error bars indicate the 95% confidence interval. **(b).** Illustrations of the Allen CCFv3 with isocortex structures colored by the relative plaque density of their corresponding module in each mouse line.

To assist in visualizing the whole brain, 3D, regional distribution patterns of methoxy-X04 labeled plaques for the different ages and mouse lines, we created 3D heatmaps (**Figure 9, Figure 10**). These maps are based on the Allen CCFv3 with each annotated brain structure colored by plaque density. **Figures 9 and 10** show example images from different 3D locations (coronal, sagittal, and horizontal slices) in 13-month-old APP/PS1 and hAPP-J20 maps, and 19-month-old Tg2576 maps (since plaque density was very low at 13 months in this line). **Movie 1** shows a flythrough of these three maps in the coronal plane. Maps for all mouse line/age group combinations are available for download at http://download.alleninstitute.org/publications/ and can be viewed in image software such as ITK-SNAP (http://www.itksnap.org/). The data and code for producing plaque maps is also available at https://github.com/AllenInstitute/plaque_map.

**Figure 9.**
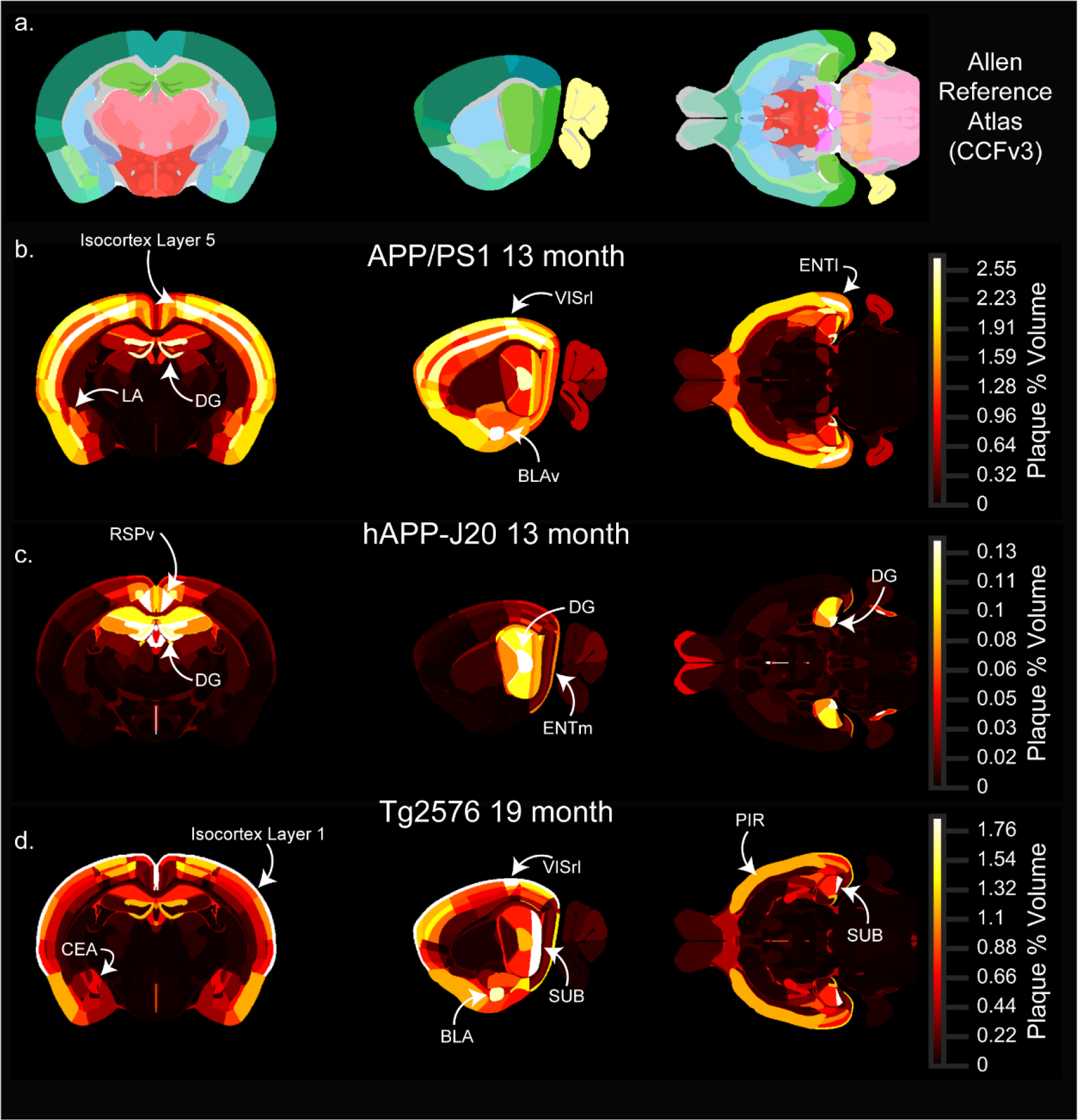
Whole brain 3D heatmaps showing the anatomical distribution of plaques. Single coronal (left), sagittal (middle), and horizontal (right) planar views from the associated full 3D map for each age-mouse line combination. **(a)** Sections from the Allen CCFv3 reference atlas showing structure annotation at the summary structure level. **(b-d)** Single sections corresponding to the reference atlas plates in the top row colored by plaque density. APP/PS1 **(b)** and hAPP-J20 **(c)** maps for 13-month-old animals are shown, and the Tg2576 **(d)** map is for 19-month-old animals. Each plaque map has its own scaled colormap (indicated in color bar on right) since absolute plaque levels are variable between ages and mouse lines. The full 3D plaque maps for all ages and mouse lines are available to download as nrrd files online at [link to be provided]. Abbreviations: LA: Lateral amygdalar nuclus, DG: dentate gyrus, VISrl: rostrolateral visual area, BLAv: Basolateral amygdalar nucleus, ventral part, ENTl: lateral entorhinal cortex, RSPv: ventral retrosplenial cortex, ENTm: medial entorhinal cortex, CEA: Central amygdalar nucleus, SUB: subiculum, BLA: Basolateral amygdala, PIR: Piriform cortex.

**Figure 10.**
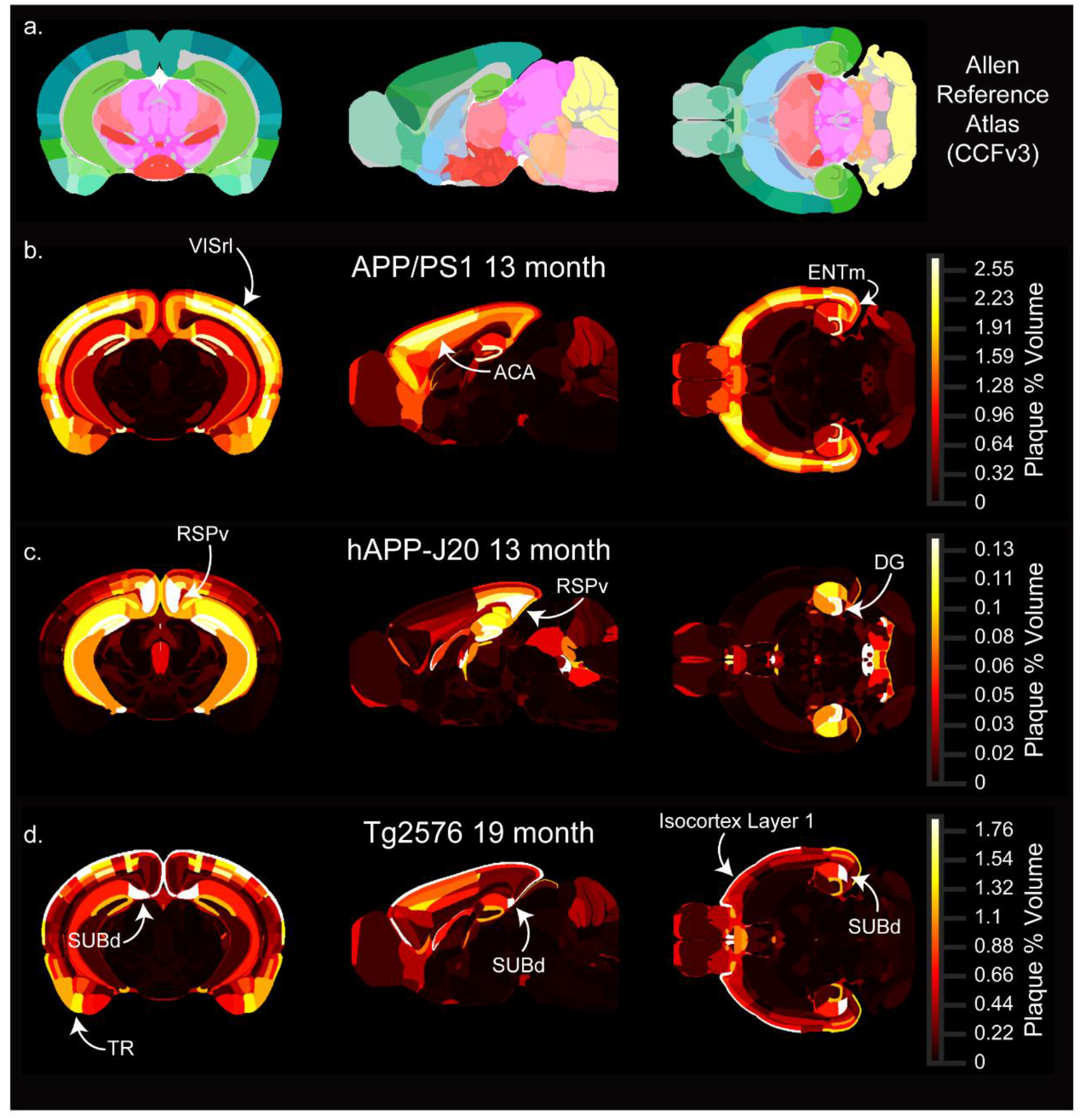
Whole brain 3D heatmaps showing the anatomical distribution of plaques at other locations. (Single coronal (left), sagittal (middle), and horizontal (right) planar views from the associated full 3D map for each age-mouse line combination. **(a)** Sections from the Allen CCFv3 reference atlas showing structure annotation at the summary structure level. **(b-d)** Single sections corresponding to the reference atlas plates in the top row colored by plaque density. APP/PS1 **(b)** and hAPP-J20 **(c)** maps for 13-month-old animals are shown, and the Tg2576 **(d)** map is for 19-month-old animals. Each plaque map has its own scaled colormap (indicated in color bar on right) since absolute plaque levels are variable between ages and mouse lines. The full 3D plaque maps for all ages and mouse lines are available to download as nrrd files online at [link to be provided]. Abbreviations: VISrl: rostrolateral visual area, ACA: anterior cingulate cortex, ENTm: medial entorhinal cortex, RSPv: ventral retrosplenial cortex, DG: dentate gyrus, SUBd: dorsal subiculum, TR: Postpiriform transition area.

One spatial feature of amyloid deposition that is easily visualized using the plaque heatmaps is the distribution of plaques by layer in the isocortex. The high density of plaques in layer 5 for APP/PS1 mice and layer 1 for Tg2576 mice are prominent features in the 3D maps (i.e. compare **Figure 9b** with **Figure 9d**). To quantify differences in plaque density across cortical layers, we calculated the plaque density per layer across all isocortical structures and compared it to the plaque density that would be expected with even plaque distribution based on the relative volume of each layer in the Allen CCFv3. The relative plaque density in each layer for the three mouse lines is plotted in **Figure 11**. Plaque density in APP/PS1 mice was more uniform across layers than the other two lines, but was higher in layer 5 (p<0.0001 than in layers 4, 6a, and 6b; p=0.03 vs. layer 1; 2-way ANOVA with Sidak’s multiple comparisons test) (**Figure 11a,b**). There appeared to be some regions such as VISrl where plaques were as dense in layer 2/3 as in layer 5 (**Figure 9b, Figure 10b**), and indeed the density in layer 5 was not different from layer 2/3 (p=0.8, 2-way ANOVA with Sidak’s multiple comparisons test). In the retrosplenial cortex of hAPP-J20 mice, plaque density was highest in layer 5 **(Figure 9c, Figure 10c)**, but in other cortical regions, hAPP-J20 and Tg2576 mice both had the highest plaque density in layer 1 (**Figure 11c-f, Figure 9c,d; Figure 10c,d**). Across all regions, both Tg2576 and hAPP-J20 mice had higher plaque density in layer 1 than in any other layer (p<0.0001, 2-way ANOVA with Sidak’s multiple comparisons test). The strong preference for layer 1 plaques in Tg2576 and hAPP-J20 mice appeared to come partly from the surface vasculature which had prominent methoxy-X04 labeling in Tg2576 mice at all ages and 19-month-old hAPP-J20 mice (**Figure 4**). In Tg2576 mice, there was no difference in the relative plaque density between layer 2/3 and layer 5, but hAPP-J20 mice had significantly higher plaque density in layer 5 than in layer 2/3 across the whole cortex (p=0.007, two-way ANOVA with Sidak’s multiple comparisons).

**Figure 11.**
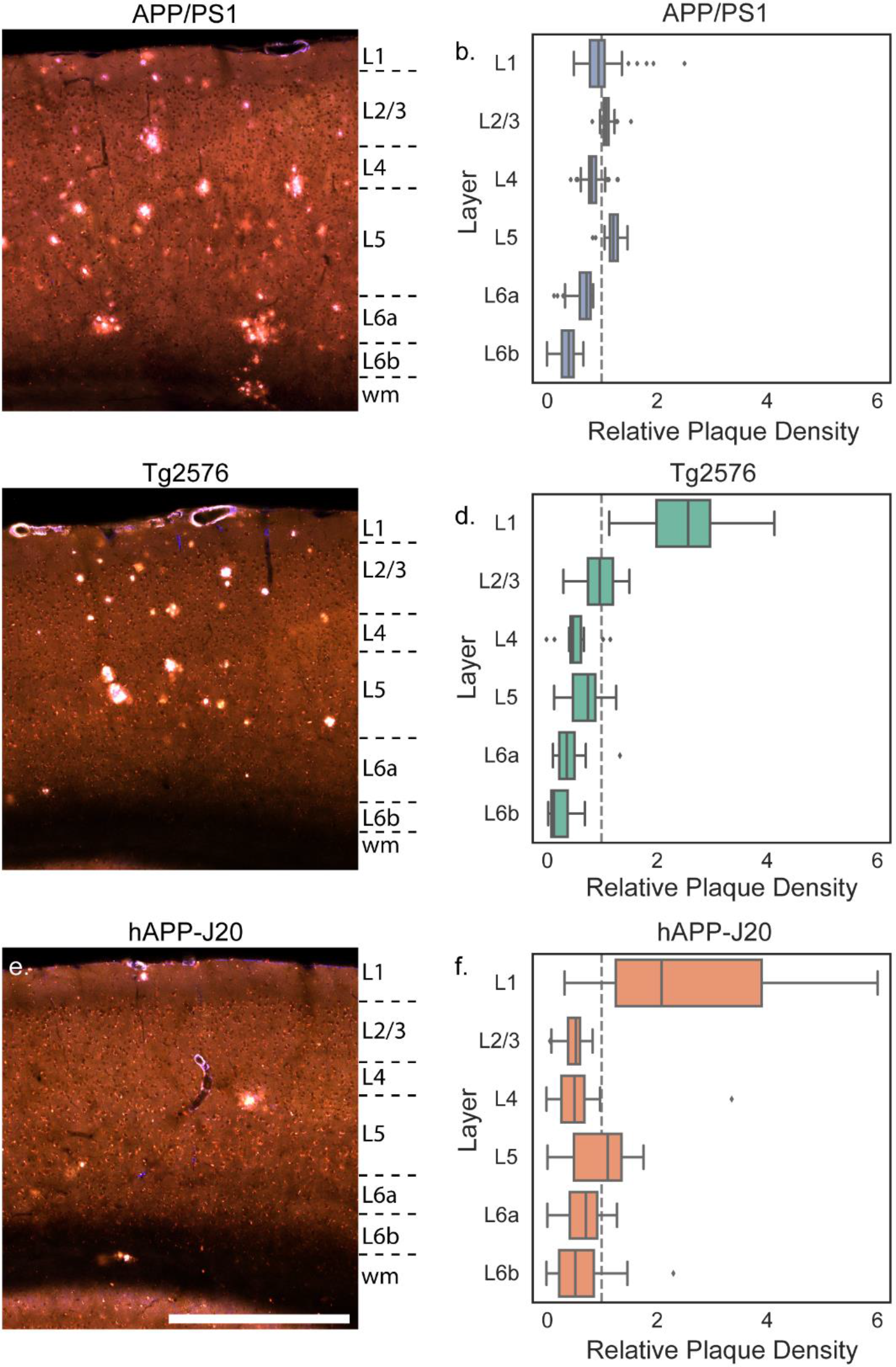
Plaque distribution across cortical layers differs between mouse lines. Images showing plaques in the parietal cortex of 19-month-old APP/PS1 **(a)**, Tg2576 **(b)**, and hAPP-J20 **(c)** mice. Approximate layer boundaries are indicated in text to the right of each image (wm = white matter). Scale = 500 μm. The relative plaque density in each cortical layer across the entire isocortex is plotted separately for APP/PS1 **(b)**, Tg2576 **(d)**, and hAPP-J20 **(f)** mouse lines. Box plots show median and IQR with whiskers extending up to 1.5 times the IQR. Outliers are plotted as individual points.

In summary, we quantified patterns of plaque deposition across brain regions at the major division and summary structure level for three APP transgenic mouse lines. We found line-specific differences in plaque density for different brain divisions and for structures within major brain divisions. The pattern of plaque deposition in the isocortex differed between mouse lines at both the regional and layer levels.

### 3.5 Comparison of plaque deposition patterns between mouse lines and human

In AD patients, plaque accumulation follows a predictable pattern starting in the isocortex, followed by hippocampal regions, then other subcortical regions such as striatum, basal forebrain, and thalamus, and finally brainstem nuclei and cerebellum. These stages of Aβ deposition were described by (Thal, Rüb, et al., 2002) based on quantifying the percent of human tissue samples with plaques in a list of regions across the whole brain. To compare the brain-wide patterns of plaque deposition described here with the pattern of Aβ deposits observed in autopsy tissue from patients in different phases of AD, we examined plaque density in rodent homologs to the human brain regions used to quantify AD staging in (Thal, Rüb, et al., 2002). For each region reported to contain Aβ deposits (left column, **Figure 12a**), the closest mouse homolog from the Allen CCFv3 reference atlas is listed in the second column. In cases where there was more than one mouse structure that could correspond to the human brain region listed, we used the median plaque density across all the corresponding structures. We calculated the median plaque density per structure across ages for each of the three APP mouse lines and, for APP/PS1 and hAPP-J20 mouse lines, we subtracted the median false positive value for each structure in control mice of the same line. We applied a color map spanning from the 10^th^ to 90^th^ percentile of plaque densities for **all** annotated structures (839) for each group to create a table similar to the one in (Thal, Rüb, et al., 2002) (**Figure 12a**). The temporal and regional pattern of plaque deposition for APP/PS1 and Tg2576 mice was remarkably similar to the pattern reported by Thal et al. with a few notable exceptions. Most importantly, plaques accumulated in the cerebellar cortex (CBX) very early for APP/PS1 mice and were also seen in Tg2576 mice, but plaques were not regularly seen in the cerebellum (molecular layer or granule cell layer) until the last phase of Aβ deposition, Phase 5, in human brain tissue. Plaque density was high in central gray for all three mouse lines and in the substantia nigra for APP/PS1 and hAPP-J20 mouse lines, but these were both regions in which we observed false positive signal in our automated segmentation, so these values may overestimate the true plaque signal even though we subtracted the control data values. In the top portion of the chart in **Figure 12a**, the plaque distribution in APP/PS1 mice and Tg2576 mice follows a similar step-wise pattern to that described in autopsy tissue for Phase 1, Phase 2, and the beginning of Phase 3. The thalamus, hypothalamus, and striatum were key regions that defined Phase 3 for human brains. While APP/PS1 mice began to accumulate plaques in the thalamus and striatum at 13 months, at 19-months-old they did not have plaques in the hypothalamus and still had low plaque density in the basal forebrain and brainstem nuclei. This implies that even after aging for 1.5 years, APP/PS1 mice, which had the most aggressive plaque deposition of the three tested, only partially resembled human Phase 3 pathology. Tg2576 and hAPP-J20 mice had low but measurable plaque density in the thalamus, hypothalamus, and striatum, but we did not observe plaques in these regions so these values were likely false positives.

**Figure 12.**
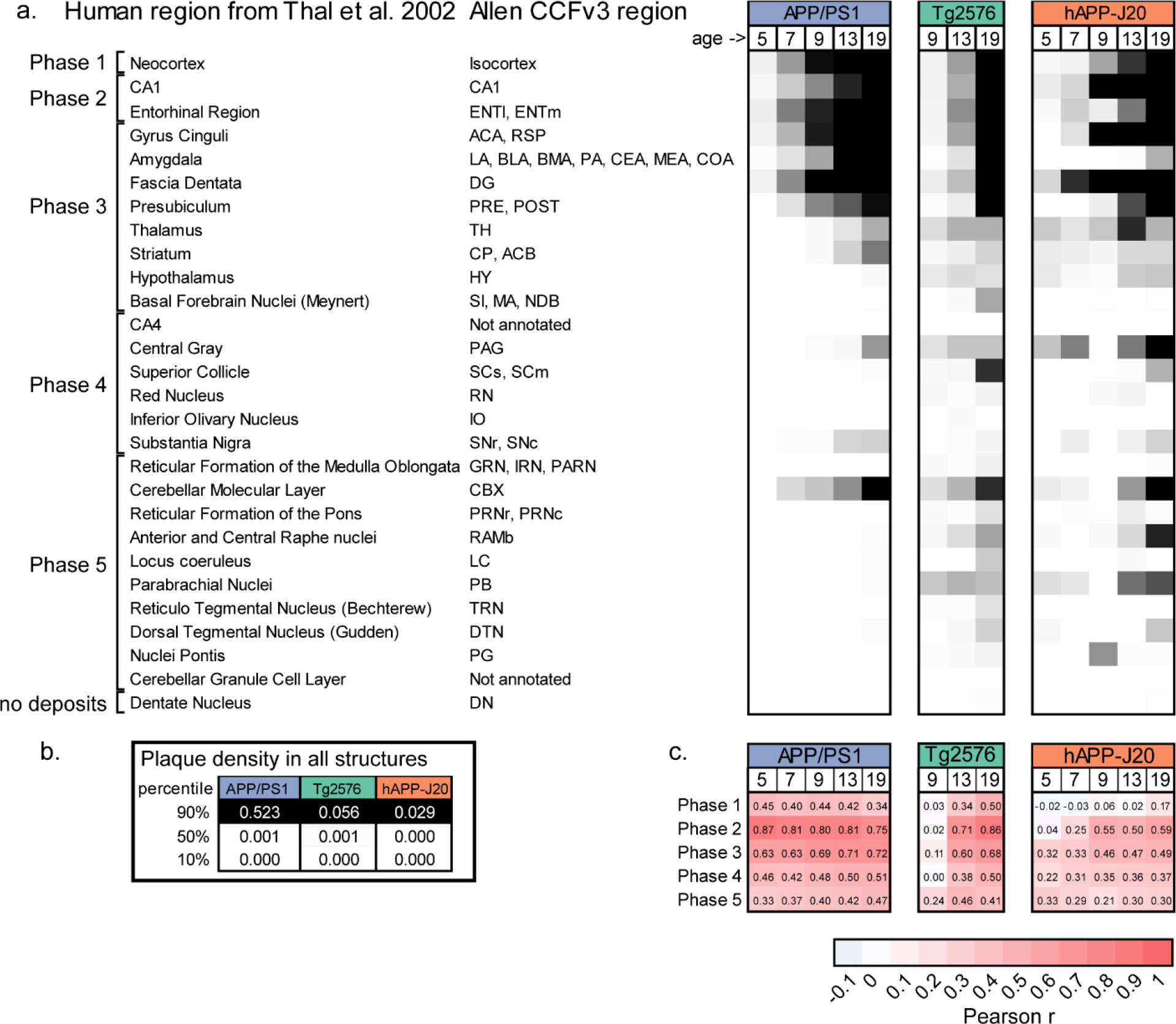
Brain-wide spatial-temporal patterns of plaque deposition in APP/PS1 and Tg2576 mice is most similar to the pattern seen in human patients with AD progression. **(a)** Comparison of relative plaque density in similar structures between human autopsy tissue and APP mouse models. Human brain regions where Aβ pathology was quantified by Thal et al. (2012) are listed in the left column and the corresponding region(s) from the Allen CCFv3 reference atlas are listed in the second column. Plaque density (or median plaque density where there are multiple regions) for each age group and mouse line is indicated by the heatmap in the columns to the right. The colormap spans from the 10th percentile to the 90^th^ percentile of the plaque density for **all** structures at all ages in that mouse line. **(b)** 10%, 50%, and 95% values for each mouse line. **(c)** Similarity measured using the Pearson correlation coefficient for comparisons between plaque density in each mouse line and age group with the fraction of patients showing plaques in each region during the five Phases of Aβ deposition in patients. Abbreviations: ENTl: Entorhinal area, lateral part; ENTm: Entorhinal area, medial part, dorsal zone; ACA: Anterior cingulate area; RSP: Retrosplenial area; LA: Lateral Amygdalar nucleus; BLA: Basolateral amygdalar nucleus; BMA: Basomedial amygdalar nucleus; PA: Posterior amygdalar nucleus; CEA: Central amygdalar nucleus; MEA: Medial amygdalar nucleus; COA: Cortical amygdalar area; DG: Dentate gyrus; PRE: Presubiculum; POST: Postsubiculum; TH: Thalamus; CP: Caudoputamen; ACB: Nucleus accumbens; HY: Hypothalamus; SI: Substantia innominata; MA: Magnocellular nucleus; NDB: Diagonal band nucleus; PAG: Periaqueductal gray; SCs: Superior colliculus, sensory related; SCm: Superior colliculus, motor related, RN: Red nucleus; IO: Inferior olivary complex; Substantia nigra, reticular part; SNc: Substantia nigra, compact part; GRN: Gigantocellular reticular nucleus; IRN: Intermediate reticular nucleus; PARN: Parvicellular reticular nucleus; CBX: Cerebellar cortex; PRNr: Pontine reticular nucleus; PRNc: Pontine reticular nucleus, caudal part; RAmb: Midbrain raphe nuclei; LC: Locus ceruleus; PB: Parabrachial nucleus; DN: Dentate nucleus.

Counting the number of cases in which plaques were observed in all regions from the table would be difficult to perform computationally with our data because of the need to choose a threshold for calling a region “positive” for the presence of plaques. We instead made the assumption that plaque density and percent prevalence are related, and compared the plaque density values from our quantification with the percent prevalence values reported in (Thal, Rüb, et al., 2002). To determine which Phase of Aβ deposition was most closely matched by each experimental mouse model, we created a correlation matrix by calculating the Pearson correlation between the plaque density in all the structures in the table for each mouse line/age combination and the percent prevalence of plaques in each structure as reported by (Thal, Rüb, et al., 2002) (**Figure 12c**). For the APP/PS1 mouse line, plaque distribution was most highly correlated with Phase 2 of Aβ deposition at all ages, but the correlation between Phase 2 and Phase 3 was approximately equal for the 19-month age group confirming that 19-month-old APP/PS1 mice were transitioning to the equivalent of Phase 3. Thirteen and nineteen-month-old Tg2576 mice also had the highest correlation with Phase 2. hAPP-J20 mice, however, did not have a high correlation with any of the phases described in human cases. At 9, 13, and 19-months-old, hAPP-J20 mice had a moderate correlation with Phase 1 in human tissue (which is defined only by plaques in the neocortex), but this correlation coefficient was still low (0.5 – 0.59). In the cases used to develop the phasing system, nondemented patients had Aβ pathology in Phases 1, 2, and 3, while clinically proven Aβ cases exhibited Aβ-Phases 3, 4, and 5 (Thal, Rüb, et al., 2002). Based on this metric, 19-month-old APP/PS1 and Tg2576 mice most closely resemble patients around the time of first diagnosis.

## 4. Discussion

We adapted the imaging, automated detection, and registration pipelines previously used for brain-wide quantification of axonal projections (Oh et al., 2014) to measure whole-brain amyloid pathology in three commonly used mouse models of AD. Automated segmentation of methoxy-X04 labeled plaques, combined with the 3D registration of whole brain image series to the Allen CCF volumetric reference atlas allow us to systematically quantify brain-wide distribution of dense-core plaque pathology at high resolution for a large sample size. The processing of non-transgenic APP^−/−^ littermates with the same pipeline facilitated interpretation of results by providing a confidence metric for the automated brain-wide plaque densities reported here. We found that plaque deposition patterns were heterogeneous across structures in all three transgenic mouse lines examined, pointing to the need for robust whole brain characterization to optimize the usage of existing and novel mouse models.

Comparing estimates of plaque density across different studies and brain regions is challenging, as it requires considering the labeling method, staining conditions, and imaging conditions used in each study in addition to animal age and transgenic line. Our pipeline approach provided consistency in labeling method and imaging conditions, and our use of automatic segmentation ensured consistent detection of plaque density across many images and samples, allowing us to quantitatively compare plaque density across the whole brain with unprecedented anatomical specificity. However, quantitatively comparing results obtained with different methods of plaque labeling is still an important challenge. Here, we found that a lower fraction of total plaque area was detected with methoxy-X04 in hAPP-J20 mice compared with APP/PS1 and Tg2576 lines. We measured the magnitude of this difference by comparing algorithmically-detected methoxy-X04 plaques with antibody labeled plaques in the same sections, but it is important to note that our automatic segmentation is based on single two-photon section images while the same sections were imaged with a widefield microscope after antibody labeling. However, the ratio between the two labeling methods would not be expected to differ between mouse lines. Importantly, the fraction of individual plaque area covered by methoxy-X04 vs. Aβ antibody measured from confocal sections is similar to the ratio between methoxy-X04 and antibody labeled areas measured in large ROIs (**Figure 4f,g**), indicating that the difference in detection likely reflects a biological difference in the composition of plaques for the different mouse lines. To our knowledge, this difference in the ratio of dense-core/diffuse plaques in hAPP-J20 mice compared with other APP models has not been previously reported. It will be important to confirm this result using other methods of labeling dense-core and diffuse plaques. Critically, although our experiments underestimate total plaque density, they do represent brain-wide plaque location more precisely than any other study to date in mouse models of AD.

Despite the challenges in directly comparing results between labs, we did compare our quantified plaque density values for the isocortex and hippocampus to published measurements when possible to both establish general trends in our data as compared to others, and to evaluate our automated segmentation technique. One recent comparable study quantified Thioflavin S-labeled plaque load in the cortex and hippocampus for both APP/PS1 and Tg2576 mice (Liu et al., 2017). Our measured plaque densities are on average 35% of those reported by Liu et al. (e.g., for our 13-month-old APP/PS1 mice and Liu et al.’s 10-13-month age group: cortex median = 0.80% vs. 2.1% and hippocampus median = 0.64% vs. 1.4%; for our 19-month-old Tg2576 and Liu et al.’s 14-17-month age group: cortex median = 0.20% vs. 1.9% and hippocampal median = 0.19% vs. 0.54%). On the other hand, our plaque densities are slightly higher than another report (Garcia-Alloza et al., 2006) that also used Thioflavin S labeling to quantify plaque density in the cortex of APP/PS1 mice (for our 9-month-old age group and Garcia-Alloza et al.’s 10-month-old age group: cortex median = 0.96 (IQR 0.67-1.8) vs. ~0.13-0.22%). We also report lower densities (0.2% (7 mo), 0.5% (9 mo) vs. ~2% (7-8.5 mo)) than a group using light sheet microscopy to image methoxy-X04-labeled plaques in cleared APP/PS1 mouse brains (Jährling et al., 2015). The plaque density we measured in 9-month-old hAPP-J20 hippocampus is nearly identical to a recent report using Thioflavin S staining (0.03%, Zhang et al., 2017). Overall, it seems that the densities we measured are within reported range of previous studies, albeit on the lower end of ranges.

Previous studies have reported plaque density in the cortex and hippocampus for different AD mouse models across ages, but generally regional distribution of plaque burden is not resolved beyond these broad divisions, much less across the entire brain. Recently, a 3D survey of Alzheimer’s pathologies in transgenic mouse models including APP/PS1 and Tg2576 was reported using the iDISCO clearing method (Liebmann et al., 2016). The authors imaged plaques brain-wide using tissue clearing followed by light sheet microscopy and registration of the whole brain images to the Allen CCFv3 reference atlas. The values reported by Liebmann et al. for plaque count (plaques per mm3) are higher than ours by a factor of ~10, which could potentially reflect the higher axial resolution of light sheet microscopy (2.5 μm vs. our 100 μm sectioning interval), but may also be caused by our automated detection algorithm merging individual plaques (see **Figure 4**). Liebmann et al. did not report plaque densities for all structures across the brain, but the twelve structures with highest plaque burden in APP/PS1 mice in their study generally agreed with our findings. In the isocortex, they found the highest plaque load in layers 2/3 and 5 of primary motor cortex (MOp), layers 2/3 and 5 of barrel cortex (SSp-bfd), supplementary somatosensory cortex (SSs), and dorsal auditory cortex (AUDd). While none of these structures had significantly higher density compared to the median for the isocortex in our data, all regions were in the top 5% of densities for both our 13-month-old and 19-month-old mice. Our results were similarly in agreement with the structures in the hippocampal formation found to have high plaque load (i.e., subiculum, lateral entorhinal cortex, CA1, and the molecular layer of the dentate gyrus, **Figure 6**). There were, however, notable differences between our datasets, as Liebmann et al. reported high plaque density in fiber tracts, specifically in the piriform area and the corpus callosum. While the piriform cortex and corpus callosum both had measurable plaque densities in our data, neither was particularly high. Liebmann et al. focused more on individual plaques than on brain-wide plaque distribution, and they also reported variability in the shape of individual amyloid deposits, including the presence of elongated structures and vascular deposits.

Our method has several advantages over the clearing method used by Liebmann et al. Most importantly, we collected image data using the STP imaging system that was also used to construct the Allen CCFv3, so the variability in our 3D registration to this reference atlas was as low as is technically possible. Our sample size is also very high for some groups, increasing confidence in our results. We put a great deal of effort into developing and validating our custom segmentation algorithm as described in the results, although we did not similarly optimize our plaque counting algorithm, which could potentially be improved with future iteration. Finally, we labeled plaques with methoxy-X04 *in vivo*, rather than post-hoc, so penetration of the plaque label into the tissue was not a concern. Tissue clearing, however, has the distinct advantage of being amenable to more types of labels, and continues to be further optimized to best combine antibody labeling and high resolution whole brain imaging methods (like STP and light sheet microscopy). It will be useful to compare whole brain quantification of Aβ antibody-labeled plaque density with the methoxy-X04 density reported here, particularly for the hAPP-J20 line since its fraction of dense-core plaque labeling is lower than the other APP lines.

The advent of PiB-PET imaging as a biomarker in AD has been a major breakthrough for the field. Not only does it allow non-invasive estimation of whole-brain plaque density and deposition patterns in humans, but patients can also be diagnosed with probable AD much earlier in the disease process and clinical trials can be more closely controlled based on biomarker measurements (Counts, Ikonomovic, Mercado, Vega, & Mufson, 2017). To best align pre-clinical research in mouse models with human biomarker data, the same PET imaging methods should be applied to mice. While STP imaging of methoxy-X04 labeling gives much higher spatial resolution than possible with PET imaging, ideally one would like to seamlessly move across scales in mice to enable better experimental testing of potential mechanistic underpinnings of disease pathology that could be extensible to humans. Unfortunately, PiB does not label Aβ deposits in some mouse models, including both APP/PS1 and Tg2576 (Klunk et al., 2005; Snellman et al., 2013), but at least one quantitative comparison between amyloid PET and methoxy-X04 labeling in different mouse lines has been performed (Brendel et al., 2015). The significance of the differential affinity of imaging probes for various forms of Aβ is still an active area of research (Schilling et al., 2016).

For many years, regional distribution patterns have been studied in relationship to tau pathology and its progression with disease stage (H Braak & Braak, 1991; Heiko Braak, Alafuzoff, Arzberger, Kretzschmar, & Del Tredici, 2006). Tau progression through brain networks has more recently been interpreted as prion-like propagation between cells (Lewis & Dickson, 2016). Not to be left behind, there has been increasing recognition of the occurrence of specific patterns of regional amyloid pathology as well in AD, starting with the discovery that Aβ plaques preferentially form in regions of the brain that are part of the resting state default mode network (DMN, R. L. Buckner, 2005). Not only do AD patients have early and heavy deposition of plaques in the DMN (Mormino et al., 2011; Palmqvist et al., 2017), the activity of this brain-wide network is also increasingly impaired as disease progresses (Brier et al., 2012; Hafkemeijer, van der Grond, & Rombouts, 2012; Jones et al., 2015; Mormino et al., 2011). These functional connectivity deficits related to the DMN are robust enough that they have been proposed as a biomarker to track disease progress, although several limitations still need to be addressed before this could be implemented (Jones, 2016). The mechanisms underlying this link between Aβ pathology and functional impairment of large-scale networks have been elusive at the cellular and behavioral level, where plaque pathology, neurodegeneration, and cognitive impairment have not been well-correlated (Nelson et al., 2012). One intriguing hypothesis is centered around the findings that Aβ deposition is related to neuronal activity (Bero et al., 2011; Cirrito et al., 2008; Kamenetz et al., 2003; Li et al., 2013). A recent study directly tested this hypothesis in Tg2576 mice by depriving the barrel cortex of sensory input, which decreased neuronal activity, and measured the effect regional plaque load (Bero et al., 2011). These authors wisely began their study by quantifying plaque load across cortical areas before using sensory deprivation to perturb the distribution of plaque levels. An increased focus on brain-wide distribution of pathology, in addition to pathology levels, should be an important component of ongoing research in rodent models of AD to best align with data from patients.

In AD patients, plaque deposition expands from the DMN and other cortical structures to hippocampus and subcortical regions. Plaque pathology in the APP transgenic lines tested here mimics these patterns of regional vulnerability in different ways. hAPP-J20 mice have a cortical plaque distribution pattern that most resembles the recent characterization of a rodent DMN homolog (Lu et al., 2012; Sforazzini et al., 2014; Stafford et al., 2014; Zerbi et al., 2015). The preferential accumulation of plaques in retrosplenial and prefrontal cortex in these mice suggests that they could be useful for testing models of regional vulnerability to amyloid deposition in the cortex. In contrast, the brain-wide sequence of plaque deposition in APP/PS1 and Tg2576 mice is more similar to the pattern seen in patients (Thal, Rüb, et al., 2002), suggesting these mouse lines may be more appropriate for studies focusing on brain-wide pathology or disease progression.

There are several possible reasons for the differences we observed in spatial patterns of Aβ deposition between APP-overexpressing transgenic mouse lines. One obvious possibility is the different promotors driving transgene expression (**Table 1**) and/or their different genomic insertion sites. A recent study found differences in expression pattern for some AD mouse models using two newly developed antibodies that can differentiate between mouse and human APP (Höfling et al., 2016). Tg2576 mice were included in their study but unfortunately APP/PS1 and hAPP-J20 mice were not. However, this approach could be applied to additional mouse lines to characterize the interaction between APP expression and Aβ deposition. Interpreting the relationship between APP expression and plaque density may be complicated by the fact that measurements of APP expression patterns are dominated by signal in cell bodies, but APP levels in axons may be more relevant to plaque deposition. Testing this hypothesis by correlating APP and plaque levels in the same region or in synaptically connected regions requires quantification of both APP expression and plaque deposition patterns with high spatial resolution, but this type of study could potentially reveal interesting mechanistic details of plaque formation.

Another possibility for the differences in spatial pattern of Aβ deposition is that each pattern reflects a separate aspect of cellular and network vulnerability. Selective vulnerability could occur at the regional level (e.g. from differences in activity across brain regions) or at the cellular level (e.g. from damage to specific neuronal projection types). Since the time course and type (dense-core vs. diffuse) of Aβ deposition differs across lines, each mouse lines may be susceptible to damage by different mechanisms. Unraveling the differences between mouse lines and cell types in their vulnerability to AD pathology is a challenging problem but these experiments have great potential for unlocking mechanisms of network neurodegeneration.

The age- and line-specific average brain-wide plaque maps described here are available for download at http://download.alleninstitute.org/publications/ as nearly raw raster data (nrrd) files that can be viewed with the free software ITK-SNAP (http://www.itksnap.org/) or loaded into Python or Matlab (The MathWorks, Inc.) as 3D arrays. Data and code for generating these maps in matrix or image form is available at https://github.com/AllenInstitute/plaque_map. We encourage researchers to use these maps for comparing amyloid distribution patterns between these transgenic models and choosing appropriate brain regions and mouse lines for hypothesis testing. These significant differences in amyloid deposition patterns between similar transgenic mouse lines underline the importance of careful consideration when choosing an AD mouse model (Jankowsky & Zheng, 2017) and importantly, indicate that appropriately chosen APP mouse models can be useful for modeling amyloid related brain-wide network degeneration.

## Author Contributions

Conceptualization: JAH, JDW. Supervision: JAH, JDW. Data Acquisition: JDW, AP, PB, AH. Data Curation: JDW, ARB, AM, KEH. Informatics Pipeline Development: LK, NG, WW. Data Analysis: JDW, ARB, JEK, NG, AP, AM. Visualization: JDW, ARB, AP, AM, KEH. The original draft was written by JDW and JAH, with input from JEK, LK, AP. All co-authors reviewed the manuscript.

---

**Movie 1.** Plaque maps for 13-month-old APP/PS1, 19-month-old Tg2576, and 13-month-old hAPP-J20 mouse lines. Each frame of the movie shows a coronal section from the Allen CCFv3 reference atlas with structures colored by plaque density (percent of structure volume occupied by plaque).

## Appendix 1. False positive values for all structures in the Allen CCFv3 reference atlas.

False positive plaque signal detected in APP^−/−^ control brains by our automated informatics pipeline. For each structure annotated in the Allen CCFv3 reference atlas, we list the structure acronym, unique structure id, plaque volume (volume of structure covered by plaque signal), structure volume (total volume of each structure in mm^3^), plaque density (plaque volume/structure volume, or fraction of structure area covered by plaque signal), plaque count (number of plaques in each structure), plaque count per mm^3^ (plaque count/structure volume). Values in this table are the mean for all control animals.

## Appendix 2. Plaque volume, density, and count for all structures in all animals.

For each animal in our dataset, this table lists the unique image series id, mouse line, sex, age at death (age), and age group (5 mo, 7 mo, 9 mo, 13 mo, 19 mo). The “control” column indicates mice from the control dataset (TRUE = APP^−/−^ control, FALSE = experimental animal). Each tab contains data for all 839 structures in the Allen CCFv3 with different tabs corresponding different metrics: plaque volume (volume of structure covered by plaque signal), plaque density (plaque volume/structure volume), plaque count (number of plaques in each structure), plaques per mm^3^ (plaque count/structure volume), and structure volume (which can vary slightly between experiments due to imaging artifacts).

## Acknowledgments

We thank Dr. Stefan Mihalas for helpful discussion and advice on statistical methods, Josh Royall for graphic design work, Dr. Song-Lin Ding for assistance correlating human and mouse anatomical regions, and the Animal Care, Transgenic Colony Management, Lab Animal Services, and imaging teams for mouse husbandry, tissue preparation, and imaging. Tg2576 mice were a generous gift from Dr. Xavier Figueroa at Nativis. Research reported in this publication was supported by the National Institute on Aging of the National Institutes of Health under Award Number R01AG047589 to J.A.H. The content is solely the responsibility of the authors and does not necessarily represent the official views of the National Institutes of Health. We thank the Allen Institute founder, Paul G. Allen, for his vision, encouragement, and support.

